# The replication checkpoint prevents fork stalling by reducing transcription

**DOI:** 10.64898/2026.10.17.759807

**Authors:** Jonathan Heuzé, Salomé Grassin, Philippe Pasero, Jérôme Poli

## Abstract

The replication checkpoint is essential for maintaining genome stability during replication stress. The prevailing view is that, without this pathway, stalled DNA replication forks lose structural integrity and collapse, preventing replication resumption and ultimately causing cell death. Here, we show that transcription is a major cause of replication stress in budding yeast checkpoint mutants. Checkpoint signaling limits replication defects by reducing RNA polymerase II and III occupancy at highly transcribed genes, particularly those encoding ribosomal proteins. Reducing transcription at these loci reduces fork stalling and the sensitivity of checkpoint mutants to hydroxyurea-induced replication stress. In human cells, transcription inhibition likewise improves replication recovery when ATR is inhibited, suggesting that this function is conserved. Together, our findings extend the canonical view of critical checkpoint functions: in addition to stabilizing stalled forks, it actively reduces RNA polymerase roadblocks that would otherwise impede their progression.

**Highlights:**

- The replication checkpoint acts upstream of fork stabilization to prevent stalling
- Replication defects is caused by highly expressed ribosomal proteins genes
- Reducing ribosomal genes transcription partially rescues checkpoint mutants
- This protective checkpoint function is evolutionarily conserved in human cells.

## Introduction

During every cell cycle, replication forks must navigate a genome containing numerous obstacles to replisome progression. In response to dNTP shortage, DNA polymerase inhibition, or DNA lesions, helicase-DNA polymerases uncoupling exposes RPA-coated ssDNA, activating the checkpoint kinases Mec1^ATR^ and Rad53^CHK1^ (Saldivar et al. 2017). In eukaryotes, both kinases are essential and form a primary barrier against tumorigenesis. The checkpoint orchestrates global responses including increased dNTP synthesis, cell-cycle arrest, and transcriptional induction of DNA damage response genes (Saxena et Zou 2022). It also acts locally at replication sites by inhibiting late origin firing (Santocanale et Diffley 1998; Shirahige et al. 1998; Zegerman et Diffley 2010; Mantiero et al. 2011; Koundrioukoff et al. 2013), modulating fork speed (Bacal et al. 2018; McClure et Diffley 2021), and maintaining functional coupling between the CMG helicase and DNA polymerases (Devbhandari et Remus 2020). In the absence of checkpoint activity, unrestrained helicase progression generates excess ssDNA, depleting RPA and causing a replication catastrophe (Toledo et al. 2013). Defective Okazaki fragment maturation can also exhaust PCNA-RFC complexes (Bertolin et al. 2025; Canal et al. 2025; Chhetri et al. 2026).

The checkpoint also actively shields stalled replication forks from collapse, a transition in which forks lose the ability to resume DNA synthesis after the stress is relieved (Zeman et Cimprich 2014). In budding yeast, this pathological process has been associated to fork reversal mediated by Rad5 (Sogo et al. 2002; Bertolin et al. 2025), uncontrolled nucleolytic degradation of nascent DNA strands by Exo1 (Segurado et Diffley 2008; Morafraile et al. 2020), the MRX complex (Delamarre et al. 2020), and the accumulation of homologous recombination factors such as the Rad51 and Rad52 recombinases (Núñez-Martín et al. 2025).

Transcription is a major endogenous obstacle to replisome progression, giving rise to transcription-replication conflicts (TRCs). The outcome of these encounters depends in part on the relative orientation of the two machineries, head-on collisions being more detrimental than co-directional encounters (García-Muse et Aguilera 2016). RNA:DNA hybrids can further impair fork progression and restart (Petermann et al. 2022; Heuzé et al. 2023). Highly transcribed loci such as tRNA genes, transcribed by RNAPIII, were among the first loci identified as replication pause site (Deshpande et Newlon 1996). A polar replication fork barrier (RFB) was also identified downstream of ribosomal DNA (rDNA) genes transcribed by RNAPI to prevent head-on collisions (Shyian et Shore 2021). In human cells, disrupted coordination between transcription and replication is a major source of oncogene-induced replication stress and genome instability during the early tumorigenesis (Macheret et Halazonetis 2018). Although TRCs could represent harmless events during an unperturbed S phase, replication stress could increase opportunities for collisions by prolonging the time forks spend traversing transcribed loci (Promonet et al. 2020).

Emerging evidence links the S phase checkpoint to the regulation of transcription. During replication stress, Mec1^ATR^ cooperates with the INO80 chromatin remodeler and the elongation factor PAF1 to promote the eviction of RNAPII from chromatin and prevent TRCs (Poli et al. 2016; Hurst et al. 2021). In human cells, ATR also controls RNAPII levels on chromatin during DNA damage and replication stress (Im et al. 2014; Landsverk et al. 2019). Yet, how the checkpoint control transcription genome-wide, and whether RNA polymerase clearance is important for replication fork progression and restart, remains unclear.

Here, we show that ribosomal protein (RP) and ribosome biogenesis (RiBi) genes are major targets of checkpoint-mediated transcriptional repression during hydroxyurea-induced replication stress in budding yeast. Reducing RP gene transcription restores fork progression despite checkpoint deficiency, indicating that transcription is a major driver of fork stalling that the checkpoint normally counteracts. Transcription inhibition likewise improves replication recovery in human cells treated with an ATR inhibitor.

## Results

### Transcription inhibition alleviates replication defects in checkpoint-deficient cells

We and others have shown that Mec1^ATR^ transiently represses RNA polymerase II (RNAPII) transcription during replication stress (Poli et al. 2016; Hurst et al. 2021; Kaur et al. 2023). Whether failure to restrain transcription contributes to the hypersensitivity of checkpoint mutants to replication stress remains unknown. To address this, we used cordycepin, a nucleoside analog that causes premature transcription termination (Holbein et al. 2009; Turner et al. 2021). Under chronic hydroxyurea (HU) exposure, cordycepin partially suppressed the growth defects of *mec1Δ sml1Δ* and *rad53Δ sml1Δ* mutants (Fig. 1A). We then asked whether this rescue reflected improved replication fork progression, which depends on checkpoint signaling under replication stress (Tercero et Diffley 2001; Núñez-Martín et al. 2025). DNA combing revealed that cordycepin had no effect on wild-type (WT) and *sml1Δ* strains, with a median BrdU track length of ∼ 25 kb (Fig. 1B, C). By contrast, median tracks in *mec1Δ sml1Δ* and *rad53Δ sml1Δ* cells were much shorter (14.5 and 13.9 kb, respectively; Fig. 1C). Cordycepin increased these values to 19.4 kb and 18.4 kb, restoring approximately 45% of the difference relative to WT cells (Fig. 1C). These data indicate that reducing transcription improves DNA replication when the S-phase checkpoint is compromised.

**Figure 1:**
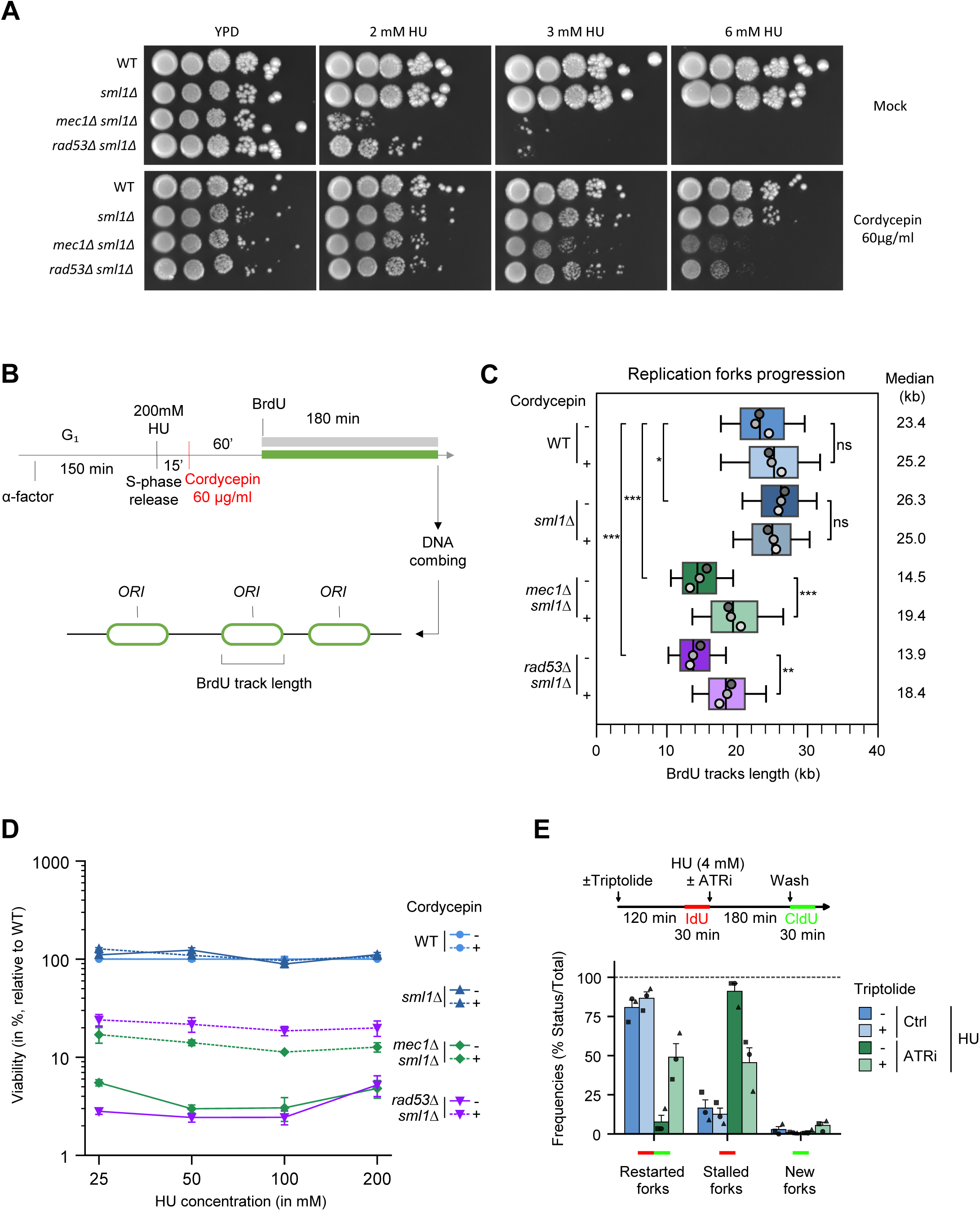
Inhibiting transcription alleviates the replication defects of checkpoint-deficient cells under replication stress. (A) Growth of cells on YPD medium +/- the indicated dose of HU +/- Cordycepin. (B) DNA combing analysis of fork progression. Exponentially growing cells were synchronized in G₁ with alpha-factor and released in S phase in the presence of 0.2M HU ± cordycepin 60µg/ml. Newly replicated DNA was labeled with BrdU for 180min after 75min in HU and analyzed by DNA combing. (C) Distribution of BrdU track lengths. Box, 25-75 percentile range. Whiskers, 10-90 percentiles range. Median is indicated in kb. One-way ANOVA performed on replicate median values followed by Tukey’s multiple-comparisons test. Significance is indicated as ns ≥ 0.05, *p <0.05, **p<0.001, ***p<0.0001. N=3 biological replicates. (D) Recovery from replication fork stalling monitored as colony outgrowth of exponentially growing cells treated with HU for the indicated concentration for 2h ± cordycepin 60µg/ml. Viability is expressed as percentage relative to the WT. Mean and SEM are indicated (N=3 biological replicates). (E) U2OS cells were treated ± Triptolide for 120min and pulsed with IdU for 30min, treated with 4 mM HU for 180min and then labeled with CldU after HU washout. Red plus green tracks: restarted forks. Red tracks: stalled forks. Green tracks: new forks (N=3 biological replicates).

We next asked whether this improvement was accompanied by increased survival after acute HU exposure. WT and *sml1Δ* cells formed colonies efficiently upon exposure to 25-200 mM HU, whereas *mec1Δ sml1Δ* and *rad53Δ sml1Δ* mutants showed severe loss of viability, with average survival of 3-4% (Fig. 1D). Cordycepin did not affect survival of the control strains (WT and *sml1Δ*), but increased survival of checkpoint mutants by approximately 30% across HU doses (Fig. 1D). Because colony formation requires cells to produce viable progeny, these data indicate that transcription inhibition improves recovery from replication stress in checkpoint-deficient cells.

To determine how cordycepin acts on checkpoint mutants, we examined chromatin occupancy of Rpb1-S2P, a marker of elongating RNAPII. ChIP-seq showed a progressive ∼20–30% reduction of chromatin-bound Rpb1-S2P at highly transcribed genes after 2-4 hours of cordycepin treatment (Fig. S1A-C). Because RNA:DNA hybrids accumulate during replication stress (Hamperl et al. 2017; Heuzé et al. 2023) and exacerbate TRCs (Uruci et al. 2026), we asked whether their removal could similarly suppress HU sensitivity. Overexpression of yeast *RNase H1* (RNH1, Fig. S1D), or bacterial *RnhA* (Fig. S1E) did not rescue checkpoint mutants, although *RnhA* complemented an *rnh1Δ rnh201Δ* control (Fig. S1F). The rescue by cordycepin is therefore associated with reduced RNAPII occupancy and was not reproduced by RNase H overexpression.

Finally, we asked whether transcription also impedes fork restart in human cells when ATR activity is inhibited following acute HU exposure. Under control conditions, 80-85% of forks restarted, irrespective of treatment with the transcription inhibitor triptolide (Fig. 1E). In contrast, ATR inhibition reduced fork restart to approximately 7%, whereas triptolide partially restored it to approximately 50% (Fig. 1E). Thus, transcription inhibition also improves fork restart in ATR-inhibited human cells, consistent with a conserved role for the replication checkpoint in coordinating transcription and replication during stress.

### Depletion of chromatin-bound RNAPII alleviates fork progression defects in checkpoint mutants

If chromatin-bound RNAPII directly impedes replication in checkpoint-deficient cells, reducing its occupancy should alleviate their replication defects. To test this independently of cordycepin, we used the anchor-away system (Haruki et al. 2008), in which rapamycin induces heterodimerization of an FRB-tagged nuclear protein with the cytoplasmic ribosomal anchor Rpl13a-FKBP12, leading to its export from the nucleus. We tagged Rpb1, the largest RNAPII subunit, and confirmed a dose-dependent reduction in its chromatin occupancy after rapamycin treatment (Fig. 2A). At 0.1 µg/ml, rapamycin reduced Rpb1 occupancy at *PYK1* and *YEF3* genes by ∼20–30%, comparable to the reduction observed with cordycepin (Fig. 2A). DNA combing revealed that this treatment had little effect on BrdU track length in WT and *sml1Δ* cells (∼23-24kb; Fig. 2B-C), but partially restored fork progression in *mec1Δ sml1Δ* (14.5kb vs 19.5kb; Fig. 2B-C) and *rad53Δ sml1Δ* cells (13.8kb vs 19.1kb; Fig. 2B-C). These results indicate that reducing chromatin-bound RNAPII restores DNA synthesis in checkpoint-deficient cells, consistent with transcription-replication conflicts contributing to their replication defects.

**Figure 2:**
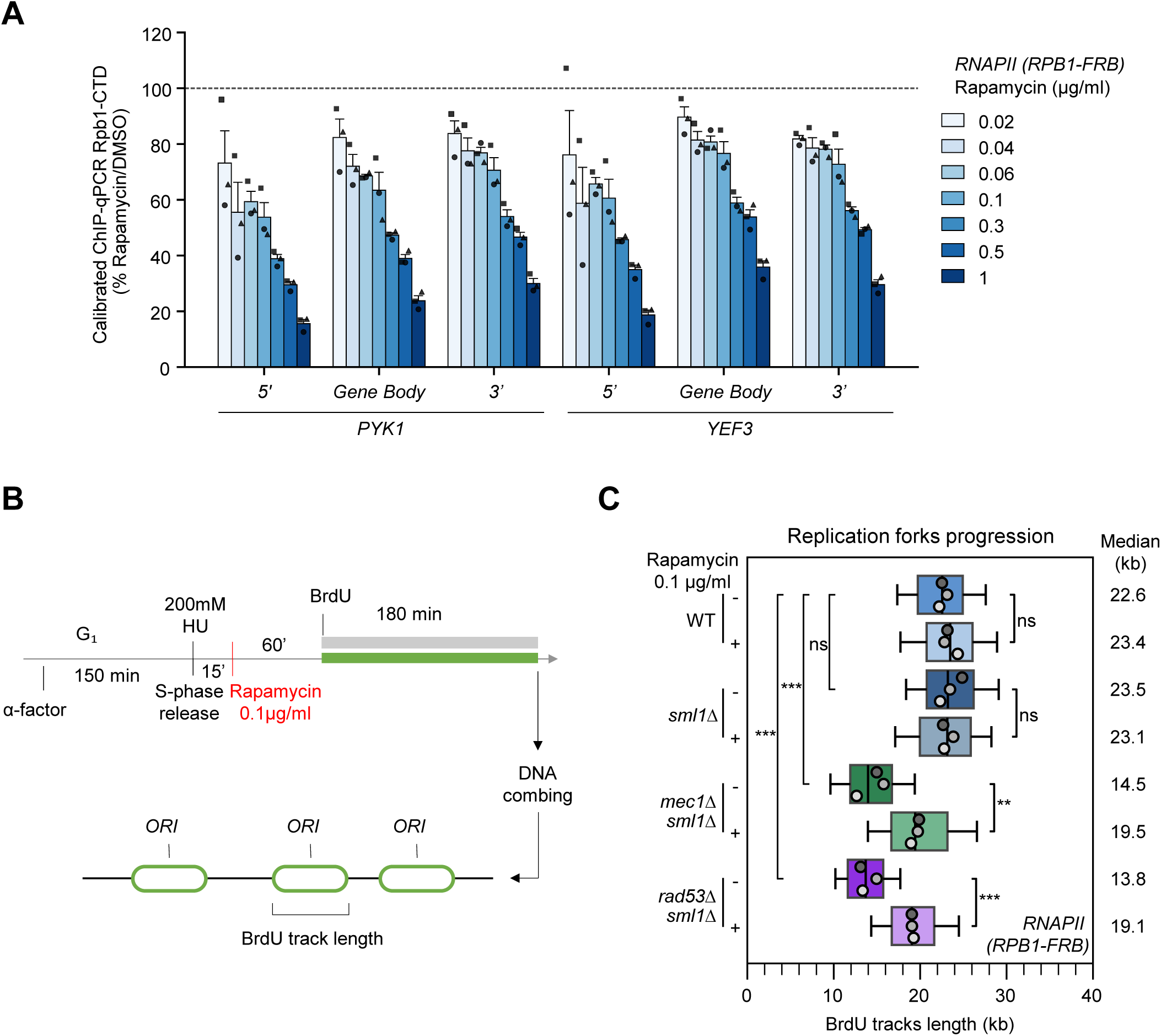
Artificial eviction of RNAPII partially rescues replication fork progression under stress in checkpoint mutants. (A) Quantitative ChIP-qPCR analysis of Rpb1-CTD enrichment following rapamycin treatment. Cells were treated with the indicated doses of rapamycin for 60 min. *RPB1-FRB* enrichment was measured across the 5’, gene body, and 3’ regions of highly transcribed genes *PYK1* and *YEF3*. Values represent %Input normalized to the average %Input of *PYK1* and *PMA1* in *C. glabrata*, displayed as a ratio relative to the DMSO condition. Data show individual biological replicates along with the mean and SEM (N = 3). (B) DNA combing analysis of fork progression. Exponentially growing cells were synchronized in G₁ with alpha-factor and released in S phase in the presence of 0.2M HU ± cordycepin 60µg/ml. Newly replicated DNA was labeled with BrdU for 180min after 75min in HU and analyzed by DNA combing. (C) Distribution of BrdU track lengths for *RPB1-FRB* background strains. Boxes represent the 25th–75th percentiles, whiskers represent the 10th–90th percentiles, and medians are indicated in kb. Statistical significance was determined using a one-way ANOVA on replicate median values followed by Tukey’s multiple-comparisons test (ns, p > 0.05; *p < 0.05; **p < 0.001; ***p < 0.0001). N = 3 biological replicates.

### Checkpoint-dependent RNAPIII eviction limits fork stalling under replication stress

In addition to RNAPII, RNAPIII can impede replication forks at tRNA genes (Deshpande et Newlon 1996). Mec1^ATR^ and Rad53^CHK1^ have been implicated in the repression of a subset of tRNA transcription during replication stress (Nguyen et al. 2010; Hurst et al. 2021), but the genome-wide extent of this regulation has not been examined. RNAPIII transcribes 275 tRNA loci and 12 non-coding RNAs in the yeast genome. To map RNAPIII occupancy quantitatively, we performed spike-in normalized ChIP-seq of Rpo31-PK, the largest subunit of RNAPIII (Fig. S2A-B and S3A). In G₁-synchronized cells, RNAPIII occupancy was broadly similar across these loci (Fig. S2C). Upon HU treatment, Rpo31 occupancy declined at nearly all loci (Fig. S2D). This reduction was partly checkpoint-dependent, *mec1Δ sml1Δ* and *rad53Δ sml1Δ* cells retaining approximately 20% more Rpo31 on chromatin than WT and *sml1Δ* controls (Fig. 3A). Metagene analysis also showed increased RNAPIII occupancy near transcription start sites (TSS) in checkpoint mutants (Fig. S2E). To test whether excess chromatin-bound RNAPIII also impedes replication fork progression in checkpoint-deficient cells, we fused Rpo31 to an FRB domain for anchor-away-mediated depletion. ChIP-qPCR confirmed that rapamycin efficiently depleted RNAPIII from chromatin (Fig. 3B). We selected a dose that decreased occupancy by 20–30% (Fig. 3B), comparable to the excess retained in checkpoint mutants. This partial depletion had no effect on BrdU track length in control strains (∼23-24kb; Fig. 3C-D), but partially alleviated the replication defects of *mec1Δ sml1Δ* (14.7kb vs 19.2kb; Fig. 3C-D) and *rad53Δ sml1Δ* mutants (14.7kb vs 19.4kb; Fig. 3C-D). These results indicate that, alongside RNAPII, chromatin-bound RNAPIII impedes oncoming replisomes under replication stress.

**Figure 3:**
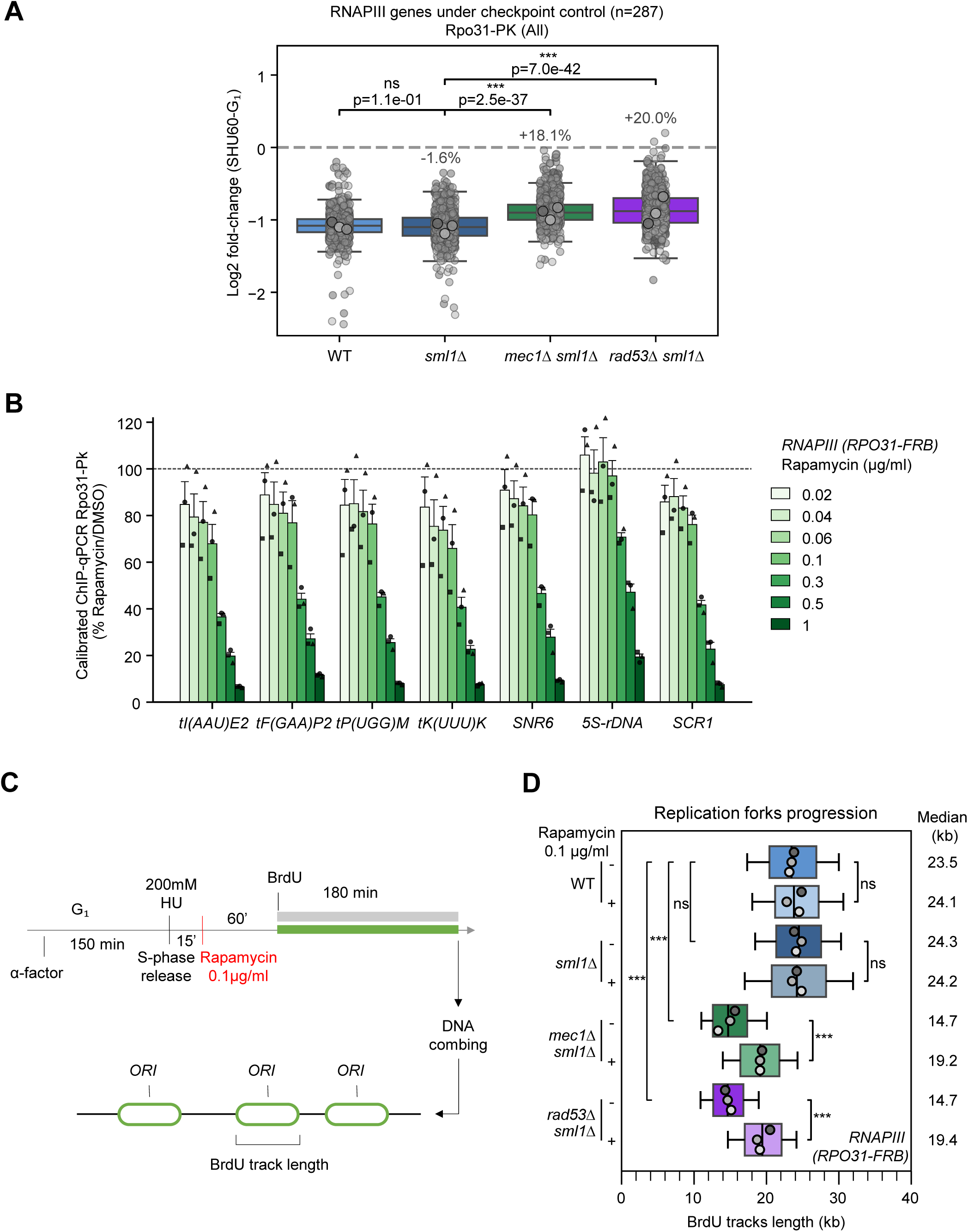
Artificial eviction of RNAPIII partially rescues replication fork progression under stress in checkpoint mutants. (A) Boxplot showing HU-induced changes in Rpo31-Pk chromatin binding across all genes (expressed as Log2-fold change [SHU60 vs G_₁_]) in mutant and control strains. Retention percentages (indicated above the boxes) were calculated from the mean Log2-fold-change differences between the indicated and control strains, normalized to the absolute mean control value (N= 3 biological replicates). Statistical significance between strains was evaluated using the Mann-Whitney *U* test on gene-wise average Log2-fold-changes (*p < 0.05; **p < 0.01; ***p < 0.001; ns, p ≥ 0.05). (B) Quantitative ChIP-qPCR analysis of Rpo31-Pk enrichment following rapamycin treatment. Cells were treated with the indicated doses of rapamycin for 60 min. Rpo31-FRB enrichment was measured across tDNA loci, *SNR6*, 5S-rDNA, and *SCR1*. Values represent %Input normalized to the average %Input of *tI(AAU)2* and *tF(GAA)2* in *C. glabrata*, displayed as a ratio relative to the DMSO condition. Data show individual biological replicates along with the mean and SEM (N = 3). (C) DNA combing analysis of fork progression. Exponentially growing cells were synchronized in G₁ with alpha-factor and released in S phase in the presence of 0.2M HU ± cordycepin 60µg/ml. Newly replicated DNA was labeled with BrdU for 180min after 75min in HU and analyzed by DNA combing. (D) Distribution of BrdU track lengths for *RPO31-FRB* background strains. Boxes represent the 25th–75th percentiles, whiskers represent the 10th–90th percentiles, and medians are indicated in kb. Statistical significance was determined using a one-way ANOVA on replicate median values followed by Tukey’s multiple-comparisons test (ns, p > 0.05; *p < 0.05; **p < 0.001; ***p < 0.0001). N = 3 biological replicates.

### The DNA replication checkpoint promotes RNAPII eviction from a subset of highly transcribed genes under replication stress

We previously showed that the checkpoint promotes Rpb1 eviction from chromatin during HU-induced replication stress (Poli et al. 2016; Hurst et al. 2021), but the genome-wide extent of this regulation remains unknown. To address this, we performed spike-in normalized ChIP-seq of total Rpb1-CTD and its elongating form, Rpb1-S2P (Fig. S3A-E). We compared G₁-synchronized cells with cells released into S phase in the presence of HU for 60 min. RNAPII occupancy declined at nearly all genes (5061/5172 genes for Rpb1-CTD; Fig. S4A and C and 5105/5172 genes for Rpb1-S2P; Fig. S4B and D). A small number of stress-responsive genes, including *RNR* genes, instead gained RNAPII occupancy (57/5172 genes for Rpb1-CTD; Fig. S4A and S5A and 26/5172 genes for Rpb1-S2P; Fig. S4B and S5B).

To account for differences in baseline occupancy, we grouped genes into low-, medium- and high-occupancy clusters based on their RNAPII signal in G₁ (Fig. S4E-G). After release into HU, RNAPII signal dropped to nearly undetectable levels in the low- and medium-occupancy clusters, whereas substantial levels persisted in the high-occupancy cluster (Fig. S6A-C). Under HU, the genome-wide Rpb1-CTD profile changed to resemble the Rpb1-S2P profile, consistent with enrichment of elongating complexes among the RNAPII that remained on chromatin (Fig. S6A). Overall, *mec1Δ sml1Δ* and *rad53Δ sml1Δ* mutants showed a decrease in chromatin-bound RNAPII similar to WT and *sml1Δ* controls (Fig. S6D). Thus, most of the RNAPII loss observed after HU treatment does not depend on the checkpoint.

Because highly transcribed genes may be prone to TRCs, we next compared RNAPII occupancy under HU in checkpoint-proficient and checkpoint-deficient cells within the high-occupancy cluster. Both checkpoint mutants retained more Rpb1 than WT cells at a subset of these genes (Fig. 4A-B). This effect was not due to *SML1* deletion, as the *sml1Δ* single mutant was indistinguishable from WT cells (Fig. 4C). More genes showed excess Rpb1 occupancy in *mec1Δ sml1Δ* (310/557, Fig. 4A) than in *rad53Δ sml1Δ* cells (220/552, Fig. 4B). The two gene sets overlapped extensively, although a subset showed excess occupancy only in *mec1Δ sml1Δ* (Fig. 4D). At genes shared between the sets, Rpb1 retention was also stronger in *mec1Δ sml1Δ* than in *rad53Δ sml1Δ* cells (Fig. 4E), consistent with a more prominent contribution of Mec1^ATR^ to RNAPII eviction during HU stress. Finally, metagene analysis of Rpb1-CTD and Rpb1-S2P at these genes showed that RNAPII accumulated toward the 3’ end of genes in checkpoint-deficient cells (Fig. S6E), consistent with the retention of elongating complexes. Together, these results distinguish two responses to HU: a widespread, largely checkpoint-independent loss of RNAPII from chromatin and a checkpoint-dependent reduction in RNAPII occupancy at a subset of highly transcribed genes. The stronger phenotype of *mec1Δ sml1Δ* cells suggests that Mec1^ATR^ contributes more prominently than Rad53^CHK1^ to the latter response, although these data do not establish whether Mec1^ATR^ acts directly or through other downstream factors.

**Figure 4:**
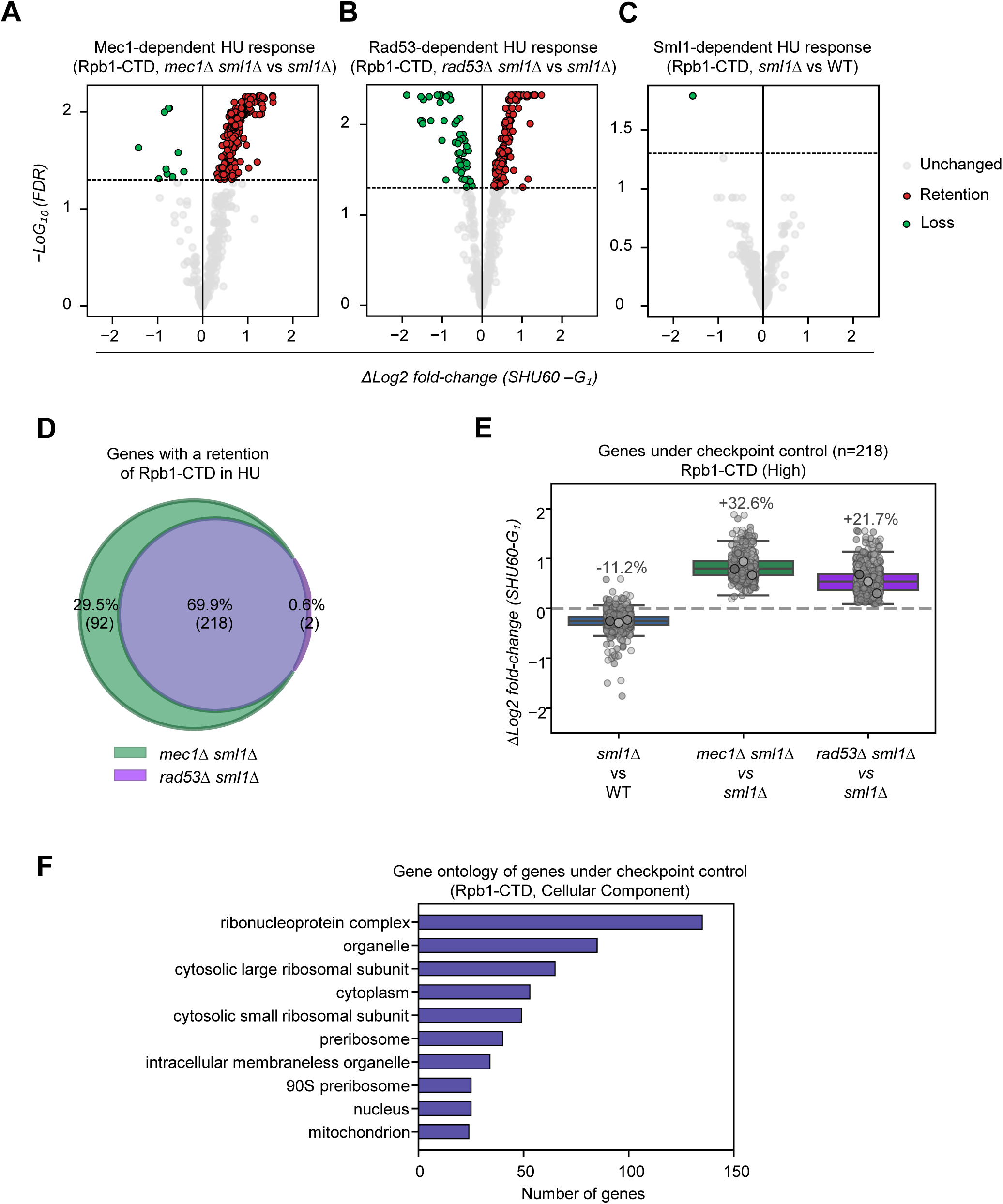
The DNA replication checkpoint drives RNAPII at a subset of highly transcribed genes under replication stress. (A-C) Volcano plots showing HU-induced changes in Rpb1-CTD chromatin binding in mutant strains relative to controls (expressed as ΔLog2-fold change [SHU60 vs G_₁_]). (A) *mec1Δ sml1Δ* versus *sml1Δ* under HU (Mec1-dependent changes): retention (n = 310), loss (n = 10), unchanged (n = 237). (B) *rad53Δ sml1Δ* versus *sml1Δ* under HU (Rad53-dependent changes)): retention (n = 220), loss (n = 54), unchanged (n = 278). (C) *sml1Δ* versus WT under HU (Sml1-dependent changes): retention (n = 0), loss (n = 1), unchanged (n = 556). In all panels, colored dots represent genes with a significantly different chromatin-binding score (adjusted p-value < 0.05; ΔLog2-fold fold change > 0 [Retention, Red] or < 0 [Loss, Green]). (D) Venn diagram showing the overlap of Mec1-dependent and Rad53-dependent genes exhibiting Rpb1-CTD retention under HU treatment. (E) Boxplot showing HU-induced changes in Rpb1-CTD chromatin binding in mutant strains relative to controls (expressed ΔLog2-fold change [SHU60 vs G₁]) for genes commonly retained in *mec1Δ sml1Δ* and *rad53Δ sml1Δ* mutants. Retention percentages (indicated above the boxes) were calculated from the mean Log2-fold-change differences normalized to the absolute mean control value. N = 3 biological replicates. (F) Gene Ontology (GO) slim mapping enrichment for Cellular Component terms among genes exhibiting Rpb1-CTD retention under HU treatment that overlap between Mec1-dependent and Rad53-dependent datasets.

### Checkpoint-dependent RNAPII eviction is specific to replication stress

The S-phase checkpoint is active at basal levels even without exogenous stress (Bastos de Oliveira et al. 2015; Forey et al. 2020). We therefore asked whether the checkpoint-dependent eviction of RNAPII observed during HU-induced stress also operates during a normal S phase. We monitored Rpb1-CTD occupancy by ChIP-seq in G₁-synchronized cells released into S phase over a 35 min time course (Fig. S7A–D). Unlike the widespread eviction triggered by HU, RNAPII occupancy showed only modest, locus-specific changes during unperturbed S phase (Fig. S8A-B). Mean Rpb1 occupancy was slightly higher in S phase than in G₁ (Fig. S8C), while α-factor-responsive genes were efficiently repressed in all S phase samples (Fig. S7E). We then compared checkpoint-proficient and checkpoint-deficient strains and found no global differences in Rpb1 occupancy (Fig. S8D). Because the checkpoint controls a subset of highly transcribed genes under HU, we specifically examined this gene set, but neither *mec1Δ sml1Δ* nor *rad53Δ sml1Δ* cells showed increased Rpb1 occupancy at these loci compared with the *sml1Δ* control (Fig. S8E). Thus, within the time course examined, we detected no checkpoint-dependent reduction in RNAPII occupancy during unperturbed S phase. This contrasts with the response to HU-induced replication stress.

### The checkpoint limits RNAPII occupancy at ribosomal protein and ribosome biogenesis genes under HU stress

The subset of highly transcribed genes whose RNAPII occupancy is controlled by the checkpoint during HU-induced replication stress may be particularly prone to TRCs. To identify features that distinguish these genes, we examined their genomic context and function (Fig. S9A). Because head-on conflicts are considered more detrimental than co-directional encounters (García-Muse et Aguilera 2016; Hamperl et al. 2017), we first assessed gene orientation relative to replication. Neither orientation was enriched among checkpoint-controlled genes (Fig. S9B). We next examined their proximity to active replication forks. In WT cells, checkpoint-controlled genes were no closer to forks as expression-matched or randomly selected genes, although highly transcribed genes as a group were closer to forks than the genome-wide baseline (Fig. S9C). Because the checkpoint restricts late origin firing (Santocanale et Diffley 1998; Zegerman et Diffley 2010; McClure et Diffley 2021), both *mec1Δ sml1Δ* and *rad53Δ sml1Δ* mutants show derepressed late origins firing upon entry into S phase in HU (Crabbé et al. 2010; Poli et al. 2012). When the resulting additional forks were included, checkpoint-controlled genes were significantly closer to active forks in checkpoint mutants, but so were expression-matched and randomly selected genes (Fig. S9D). Thus, neither orientation nor fork proximity distinguishes checkpoint-controlled genes from the comparison sets. A prominent shared feature of these genes is their high transcriptional activity. Checkpoint-controlled genes display slightly higher RNAPII occupancy than other genes within the high-occupancy cluster (Fig. S11A), while Mec1-dependent genes show intermediate RNAPII levels (Fig. S11A). Thus, although high transcriptional activity may contribute to checkpoint regulation, the relatively similar RNAPII levels across these groups suggest that it is unlikely to be the sole determinant.

To gain functional insight into these checkpoint-controlled genes, we performed a Gene Ontology (GO) analysis, which revealed a strong enrichment for translation-related functions, particularly ribosomal proteins (RP) and ribosome biogenesis (RiBi) genes (Fig. 4F, S9E). Approximately 70% of the highly transcribed genes, checkpoint-controlled genes fell into these two categories, with RP and Ribi genes accounting for 49% and (18%) respectively (Fig. 5A). Depending on the RP-gene category, the checkpoint controlled RNAPII occupancy includes between 70-95% of RP genes (Fig. S10A). In contrast, only 14% of RiBi genes belonged to the high-occupancy cluster (Fig. S10A), because most RiBi genes fell into the medium-occupancy cluster. Including the medium cluster increased the fraction of RiBi genes classified as checkpoint-controlled to 82 % (Fig. S10B-C). Gene Ontology analysis of Mec1-dependent genes showed that this subset was also enriched for translation-related functions (Fig. S11A, B), with 26 of the 92 Mec1-dependent genes classified as RP or RiBi genes (Fig. S11D). When combining genes dependent on both Mec1 and Rad53 with Mec1-dependent genes across both the high- and medium-occupancy clusters, RP and RiBi genes displayed near complete dependence on checkpoint signaling, with Mec1 making a particularly prominent contribution (Fig. S11E).

**Figure 5:**
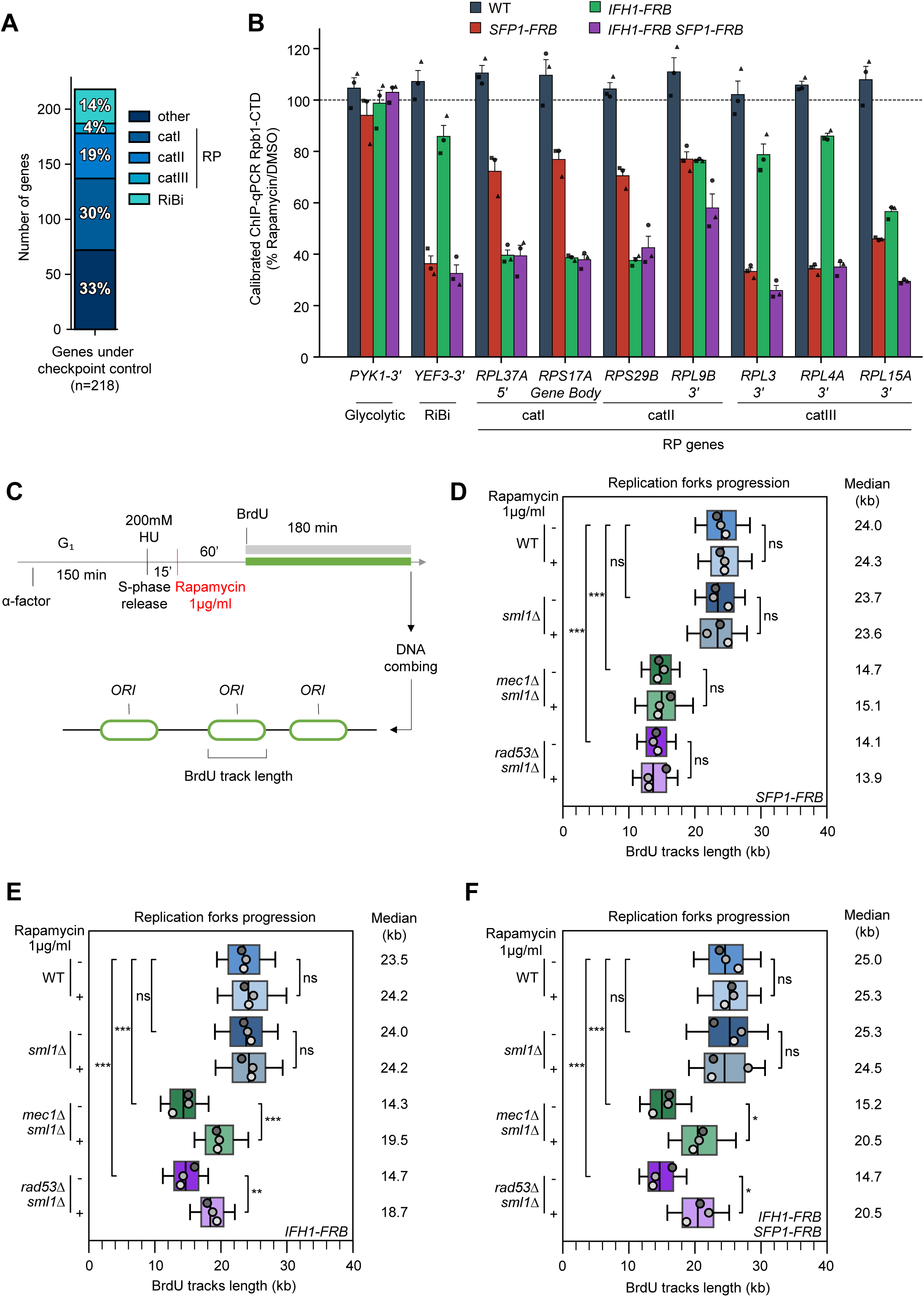
Transcription of Ribosomal Protein genes impedes DNA replication in checkpoint-deficient cells. (A) Stacked bar plot showing the distribution of RP classes and RiBi genes among genes under checkpoint control. The bar displays the absolute number of genes (y-axis) and the percentage corresponding to each class. (B) Quantitative ChIP-qPCR analysis of Rpb1-CTD enrichment. WT (FRB-untagged), *SFP1-FRB*, *IFH1-FRB*, and *IFH1-FRB SFP1-FRB* cells were treated with rapamycin at 1 µg/mL for 60 min. Values represent %Input normalized to the average %Input of *PYK1* and *PMA1* in *C. glabrata*, displayed as a ratio relative to the DMSO condition. Data show individual biological replicates along with the mean and SEM (N = 3). (C) DNA combing analysis of fork progression. Exponentially growing cells were synchronized in G₁ with alpha-factor and released in S phase in the presence of 0.2M HU ± 1µg/ml of rapamycin. Newly replicated DNA was labeled with BrdU for 180min after 75min in HU and analyzed by DNA combing. (D-F) Distribution of BrdU track lengths. Boxes represent the 25th–75th percentiles, whiskers represent the 10th–90th percentiles, and medians are indicated in kb. Statistical significance was determined using a one-way ANOVA on replicate median values followed by Tukey’s multiple-comparisons test (ns, p > 0.05; *p < 0.05; **p < 0.001; ***p < 0.0001). N = 3 biological replicates.

### Reducing RP genes expression alleviates replication defects in checkpoint mutants

RP and RiBi genes are regulated by overlapping, but distinct transcription programs. Category I and II RP genes depend primarily on Ifh1, whereas category III RP and RiBi genes depend primarily on Sfp1 ((Zencir et al. 2020; Shore et al. 2021), Fig. S10D). We therefore asked whether Ifh1 and Sfp1 contribute to the replication defects in checkpoint mutants. We first reduced RP and RiBi gene transcription by anchor-away of Ifh1 and Sfp1. First, Rpb1-CTD ChIP-qPCR confirmed that rapamycin reduced RNAPII occupancy at category I (*RPL37A* and *RPS17A*) and category II (*RPS29B* and *RPL9B*) genes upon Ifh1 depletion, and at category III (*RPL3*, *RPL4A*, and *RPL15A*) and RiBi (*YEF3*) genes (Fig. 5B) upon Sfp1 depletion, without affecting the *PYK1* control locus (Fig. 5B). *RPL9B* showed reduced RNAPII occupancy with either treatment (Fig. 5B). Depletion of Ifh1, Sfp1 or both did not impact fork progression in checkpoint-proficient control cells (WT and *sml1Δ*, Fig. 5C-F). In *rad53Δ sml1Δ* and *mec1Δ sml1Δ* cells, Sfp1 depletion alone did not improve fork progression, whereas Ifh1 depletion did (14.3 vs 19.5 kb for *mec1Δ sml1Δ*, 14.7 kb vs 18.7 kb for *rad53Δ sml1Δ*, Fig. 5C-E). This difference is consistent with the higher RNAPII occupancy at Ifh1-dependent genes relative to Sfp1-dependent-genes (Fig. S10E-F, S11F).

Sfp1 and Ifh1 have been proposed to partly compensate for one another in RP gene regulation (Shore et al. 2021). Consistent with this, co-depletion of Ifh1 and Sfp1 decreased chromatin-bound RNAPII more than either depletion alone at the loci tested (*RPL9B*, *RPL3* and *RPL15A*) without affecting *PYK1* (Fig. 5B). Co-depletion partially restored fork progression in both *rad53Δ sml1Δ* and *mec1Δ sml1Δ* mutants (15.2 vs 20.5 kb for *mec1Δ sml1Δ*, 14.7 kb vs 20.5 kb for *rad53Δ sml1Δ*, Fig. 5C and 5F) to levels comparable to those obtained after Rpb1 depletion (Fig. 2C). Together, these results indicate that reducing transcription of RP and RiBi genes is sufficient to alleviate the replication defects of checkpoint mutants, supporting a model in which checkpoint-dependent clearance of RNAPII from highly transcribed genes limits replication fork stalling under replication stress.

## Discussion

Our results identify transcription as a major contributor to the replication defects of checkpoint-deficient cells. The S-phase checkpoint has long been viewed as a pathway that stabilizes stalled forks (Tercero et Diffley 2001; Lopes et al. 2001; Cobb et al. 2003, 2005; Morafraile et al. 2015) and promotes their restart (De Piccoli et al. 2012; Cortez 2015; Gan et al. 2017). More recent work has implicated RPA exhaustion as a driver of unscheduled nuclease-mediated fork breakage (Toledo et al. 2013; Núñez-Martín et al. 2025), and PCNA depletion (Bertolin et al. 2025; Canal et al. 2025; Chhetri et al. 2026). We propose an additional upstream function of the checkpoint limiting encounters between slowed forks and RNA polymerases. Under HU treatment, checkpoint kinases promote the eviction of RNAPII from a subset of highly transcribed genes enriched in RP and RiBi functions, and contribute to the widespread reduction in RNAPIII across tRNA genes. Failure to remove either polymerase compromises replication fork progression, impairs recovery from acute HU stress, and sensitizes cells to chronic HU exposure. Reducing the chromatin occupancy of either RNA polymerase, or the transcription of RP genes, restores fork progression in checkpoint-deficient yeast cells. Transcription inhibition also restores fork restart when ATR is inhibited in human cells. By limiting transcription-associated impediments, the checkpoint could reduce the burden on its fork-protection and restart functions, although the relationship between these mechanisms remains to be tested.

Checkpoint-dependent RNAP clearance was detected under replication stress, but not during an unperturbed S phase, despite evidence of basal checkpoint kinase activity (Bastos de Oliveira et al. 2015; Lanz et al. 2018; Hurst et al. 2021). HU depletes dNTP pools and reduces fork speed from ∼ 1.5 kb/min to ∼ 0.1 kb/min (Poli et al., 2012). In HU, a fork would take roughly 15 minutes to progress through an average 1.5 kb yeast gene, but would need only ∼ 1 min under physiological conditions. Transcription occurs in discrete timed bursts rather than as a continuous process, and highly transcribed genes such as RP genes generate a productive transcript every 2 min (Pelechano et al. 2010; Lenstra et al. 2016). At normal speed, a fork would therefore have a ∼50% chance of encountering an elongating RNAPII, whereas a slow fork is expected to encounter several RNAPs. *In vitro*, multiple transcription complexes form a stronger barrier to the bacterial replisome than a single one (Brüning et Marians 2020). We therefore propose that TRCs, although rare and transient during an unchallenged S phase, become frequent at highly transcribed loci under replication stress, and that checkpoint-dependent removal of RNA polymerases limits both their frequency and severity.

Head-on collisions are generally more detrimental than co-directional ones (Azvolinsky et al. 2009; Hamperl et al. 2017; Kay et al. 2024), yet checkpoint-controlled genes showed no orientation bias. When forks are severely slowed and face several RNAPII complexes, the cumulative encounters outweigh the contribution of orientation. Likewise, RNA:DNA hybrids accumulate during replication stress and can impair fork processing and restart (Lalonde et al. 2021; Hamperl et al. 2017; Heuzé et al. 2023; Stoy et al. 2023; Audoynaud et al. 2023). R-loops have been implicated in the replication defects of mediator-deficient cells (Uruci et al. 2026). However, overexpression of RNase H1 or RnhA did not alleviate the HU hypersensitivity of checkpoint mutants, although RnhA complemented the *rnh1Δ rnh201Δ* cells. This result argues against R-loops being the main source of TRCs in these cells, although we cannot exclude a contribution from R-loops at individual loci, as we did not directly measure their abundance or removal. Together, our findings support chromatin-bound RNA polymerases as important obstacles to forks under replication stress, independently of R-loops.

Both RNAPII and RNAPIII appear to contribute to TRCs during replication stress, but their chromatin occupancy is regulated differently. Unlike the global checkpoint-mediated repression of tRNA gene transcription reported previously (Nguyen et al. 2010), the widespread decline in RNAPIII occupancy, with checkpoint mutants retaining only about 20% more than controls, indicates that checkpoint-independent mechanisms also contribute to downregulating RNAPIII during replication stress. RNAPII occupancy also fell genome-wide, but this broad reduction was largely checkpoint independent. In contrast, checkpoint activity was required to limit occupancy only at a subset of highly transcribed genes enriched in RP and RiBi genes. Global RNAPIII repression is mediated by the Rad53-dependent dephosphorylation of the Maf1 transcriptional repressor (Nguyen et al. 2010). Mec1^ATR^ was previously shown to cooperate with INO80 and PAF1C to repress transcription under replication stress (Poli et al. 2016; Hurst et al. 2021), but these general factors are unlikely to account for the locus specificity observed here. RP and RiBi genes are instead regulated by dedicated transcription factors such as Ifh1 and Sfp1, acting downstream of TORC1, raising the attractive possibility that checkpoint kinases directly target this pathway. Supporting this model, Rad53^CHK1^ interacts directly with Ifh1 through its FHA1 domain (Smolka et al. 2006). Although, this interaction is lost in the *rad53-R70A* mutant, this allele also impairs interaction with several other targets, making it difficult to isolate the specific contribution of Ifh1 (Smolka et al., 2006). Rad53^CHK1^ could therefore act as a scaffold to promote the recruitment or phosphorylation of Ifh1 by checkpoint kinases. Upon HU-induced replication stress, checkpoint-dependent phosphorylation of Ifh1 might alter its binding to Fhl1, thereby reducing Ifh1 occupancy to RP promoters and driving RP gene repression. Highlighting the critical role of this interaction, deleting the Fhl1 FHA domain, or introducing the *fhl1-G303A* or *ifh1-T681A* mutations disrupts the Ifh1–Fhl1 complex, causing a loss of Ifh1 at RP promoters and severe growth defects (Martin et al. 2004; Rudra et al. 2005; Kim et Hahn 2016). Together, these observations suggest a model in which checkpoint activation tunes RP gene expression by modulating the Ifh1–Fhl1 complex, though direct experimental validation of checkpoint-dependent phosphorylation Ifh1 phosphorylation remains to be established. Finally, Mec1^ATR^ appears to play a more prominent role than Rad53^CHK1^ in promoting RNAPII eviction, as a larger set of genes were affected and RNAPII retention was more pronounced in its absence, despite both kinases targeting a largely overlapping gene set. Whether Mec1^ATR^ acts only through Rad53^CHK1^ in this process or if it is acting on other downstream effectors remains to be determined.

Our data further suggest that the impact of TRCs depends not only on fork velocity, but also on transcriptional load. Transcription can impede fork progression and its inhibition reduces fork asymmetry (Ribeyre et al. 2016; Macheret et Halazonetis 2018; Chang et al. 2019) and transcription-associated recombination (Aguilera et Gaillard 2014). Among the transcriptional targets of the checkpoint, we found a strong enrichment in RP and RiBi genes, which are among the most heavily transcribed loci in the yeast genome. Reducing their transcription by anchor-away of Sfp1 and Ifh1 (Wade et al. 2004; Shore et al. 2021) rescued replication defects in checkpoint-deficient cells, as did cordycepin, a broad transcription inhibitor that may also dampen mTOR signaling (Lawrence et al. 2025). RNAPII occupancy was higher at RP genes than at RiBi genes. Moreover, reducing RP transcription alone (Ifh1 anchor-away) rescued fork progression whereas targeting RiBi genes (Sfp1 anchor-away) did not. These findings support the view that transcription load is a major determinant of fork stalling. Oncogene-induced hypertranscription, notably at RP genes, has likewise been shown to drive TRC formation in human cells (Kelly et al. 2026). Together, these observations suggest that replication forks are particularly vulnerable to TRCs at highly transcribed loci, especially when their progression is slowed. In human cells, ATR is critical for tolerating replication stress and restarting forks once the stress is relieved (Saxena et Zou 2022). Here, transcription inhibition with triptolide partially restored fork restart (50% vs 7% in the absence of transcription inhibition) in ATR-inhibited cells submitted to HU. This suggests that coordinating transcription with replication under stress is a conserved function of the S-phase checkpoint from yeast to humans.

Together, our findings suggest that beyond stabilizing stalled forks and promoting their restart, the S-phase checkpoint limits the impact of transcription on fork stalling, particularly at highly transcribed genes. Given that ATR is a therapeutic target in tumors experiencing high replication stress, and that oncogene-driven hypertranscription generates TRCs, it will be important to determine whether dependence on this checkpoint creates an actionable vulnerability in cancer cells.

## Supplemental Figure legends

**Figure S1:**
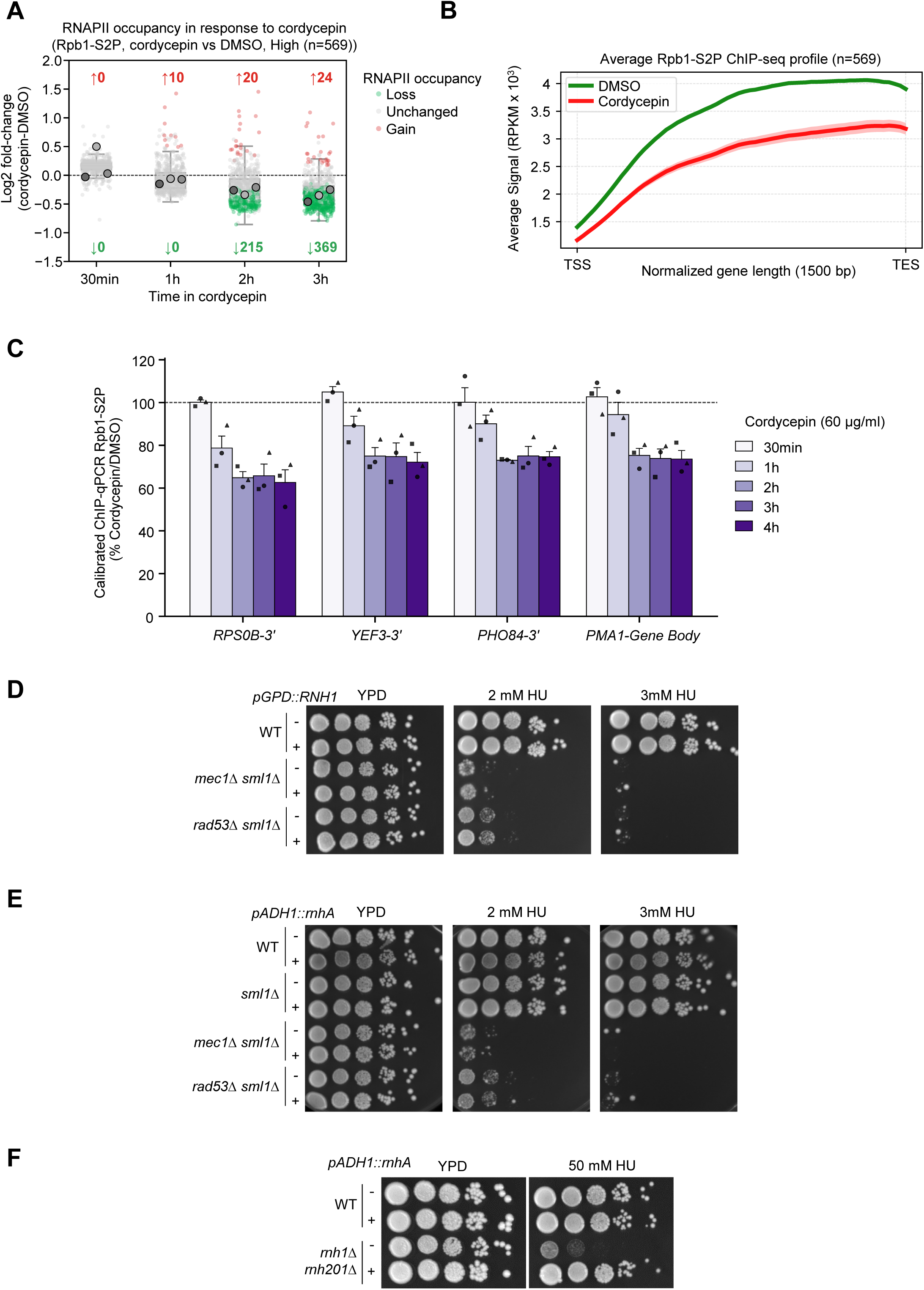
Inhibition of transcription by cordycepin alleviates the need of a functional S-phase checkpoint under HU stress. (A, B) Quantitative ChIP-seq analysis of Rpb1-S2P enrichment, in WT cells treated with 60 µg/ml of cordycepin over a time-course. (A) Boxplot showing cordycepin-induced changes in Rpb1-S2P chromatin binding relative to the DMSO control (expressed as Log2-fold change [cordycepin vs DMSO]) for highly transcribed genes. Genes exhibiting an adjusted p-value < 0.05 and a Log2-fold change > 0 are in red (Gain), while those < 0 are in green (Loss). N = 3 biological replicates. (B) Representative average profile of Rpb1-S2P along highly transcribed genes for the DMSO and cordycepin treatments at the 3 h time point. Mean and SEM are indicated (N = 3 biological replicates). (C) Quantitative ChIP-qPCR analysis of Rpb1-S2P enrichment in WT cells ± 60 µg/ml of cordycepin over a time-course. Enrichment was measured across the highly transcribed gene *PMA1*, the RP gene *RPS0B* (catI), the *RiBi* gene *YEF3*, and *PHO84*. Values represent the percentage of input (%Input) normalized to the average %Input of *PYK1* and *PMA1* in *C. glabrata*, and are displayed as a ratio relative to the untreated condition. Data show individual biological replicates along with the mean and SEM (N = 3). (D-F) Growth of indicated strains on YPD medium ± the indicated dose of HU. Spots correspond to 1:10 serial dilutions. (D-F) Growth of cells ± overexpression of pGPD(TDH3)-*RNH1*(D), pADH1-*rnhA* (*E. coli* RNase H)(E and F).

**Figure S2:**
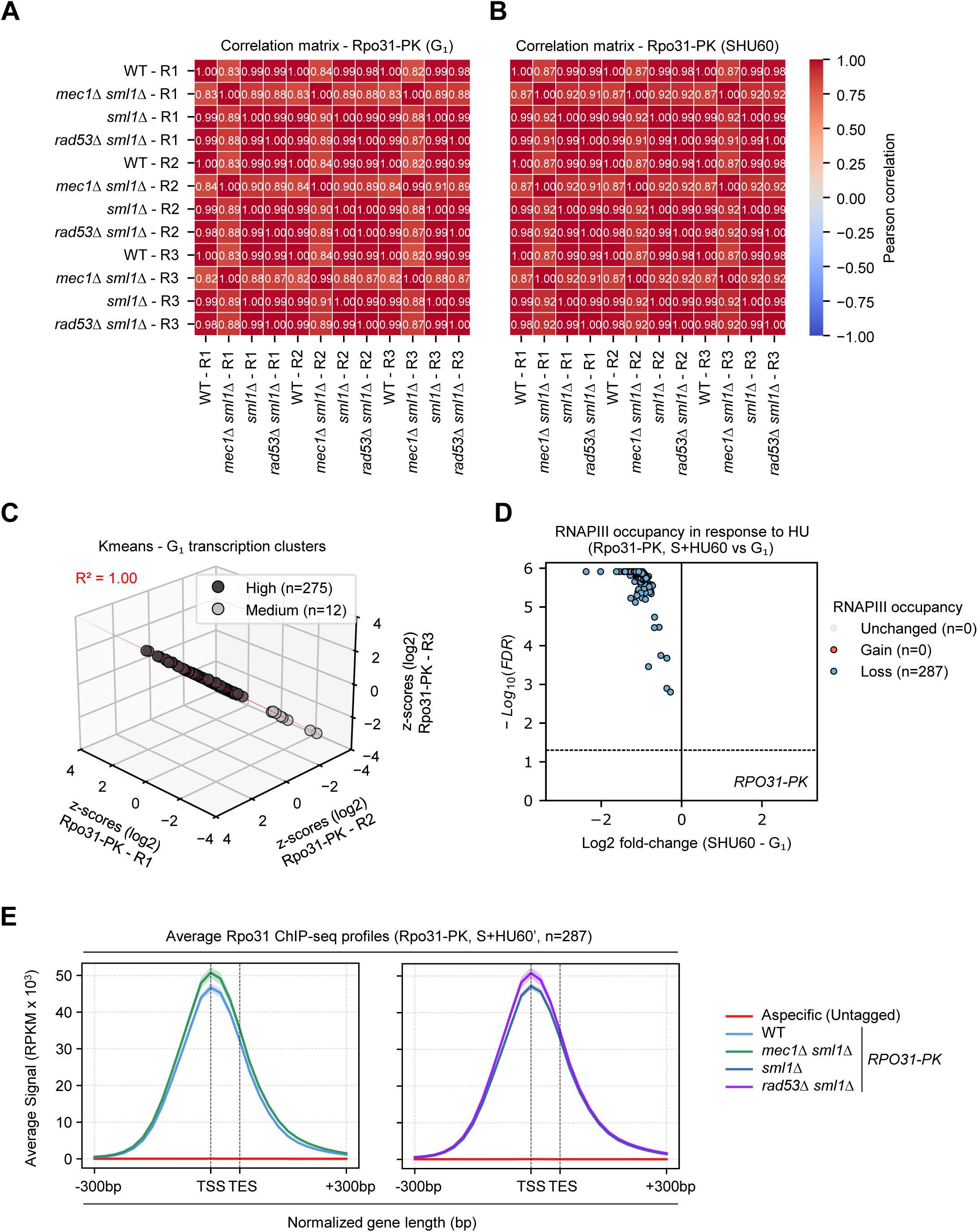
The DNA replication checkpoint drives RNAPIII eviction from chromatin under HU stress. (A, B) Pearson correlation heatmaps assessing biological reproducibility. Pairwise Pearson correlation coefficients (r) between replicate scores of Rpo31-Pk across strains and conditions. The color scale ranges from −1 to 1, indicating negative to positive correlation. (A) Rpo31-Pk in G_₁_ for 3 biological replicates, comparing all strains. (B) Rpo31-Pk in S+HU60 for 3 biological replicates, comparing all strains. (C) 3D scatter plot of G₁ transcription levels and biological replicate reproducibility (N=3) for the WT strain. Genes were categorized into three distinct clusters (High, Medium, and Low) based on Rpo31-Pk signal intensities using K-means clustering (k = 3) on log2-transformed, standardized (z-scores) RPKM total scores across biological replicates. Planar regression was applied to evaluate replicate reproducibility, and the corresponding coefficient of determination (R^2^) is shown. (D) Volcano plot showing HU-induced changes in Rpo31-Pk chromatin binding in S+HU60 compared to G₁ in the WT strain (expressed as Log2 fold change [S+HU60 vs G₁]). Colored dots represent genes with a significantly different chromatin-binding score (adjusted p-value < 0.05; Log2-fold change > 0 [Gain, Red] or < 0 [Loss, Blue]). (E) Average Rpo31 occupancy profiles from −300bp relative to the TSS to +300bp relative to the TES. Data are expressed as spike-in normalized RPKM across all RNAPIII ORFs (n = 287) plotted over a normalized gene length (75bp). Mean profiles ± SEM are shown for three biological replicates. Graphs are split into separate panels due to profile overlap between WT and *sml1Δ* strains.

**Figure S3:**
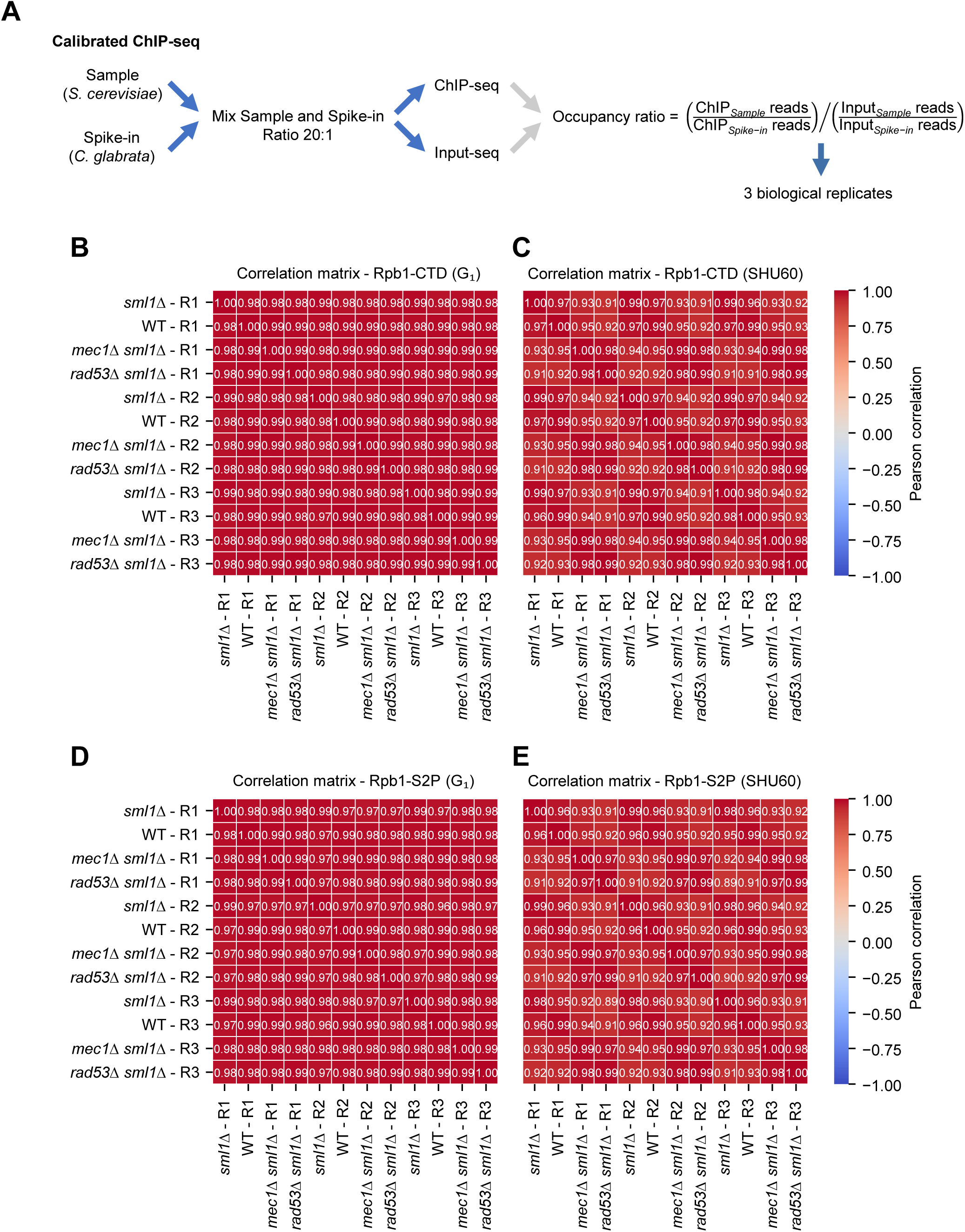
Quantitative ChIP-seq procedure and Rpb1 correlation matrices. (A) Schematic workflow of the quantitative ChIP-seq procedure (N=3). *S. cerevisiae* (sample) and *C. glabrata* (spike-in) cells were mixed prior to the ChIP assay at a 20:1 ratio. Following sequencing across three biological replicates, occupancy ratios were determined using mapped reads from both genomes. (B-E) Pearson correlation heatmaps assessing biological reproducibility. Pairwise Pearson correlation coefficients (r) between replicate scores of Rpb1-CTD and Rpb1-S2P across strains and conditions. The color scale ranges from −1 to 1, indicating negative to positive correlation. (B) Rpb1-CTD in G₁ for 3 biological replicates, comparing all strains. (C) Rpb1-CTD in S+HU60 for 3 biological replicates, comparing all strains. (D) Rpb1-S2P in G₁ for 3 biological replicates, comparing all strains. (E) Rpb1-S2P in S+HU60 for 3 biological replicates, comparing all strains.

**Figure S4:**
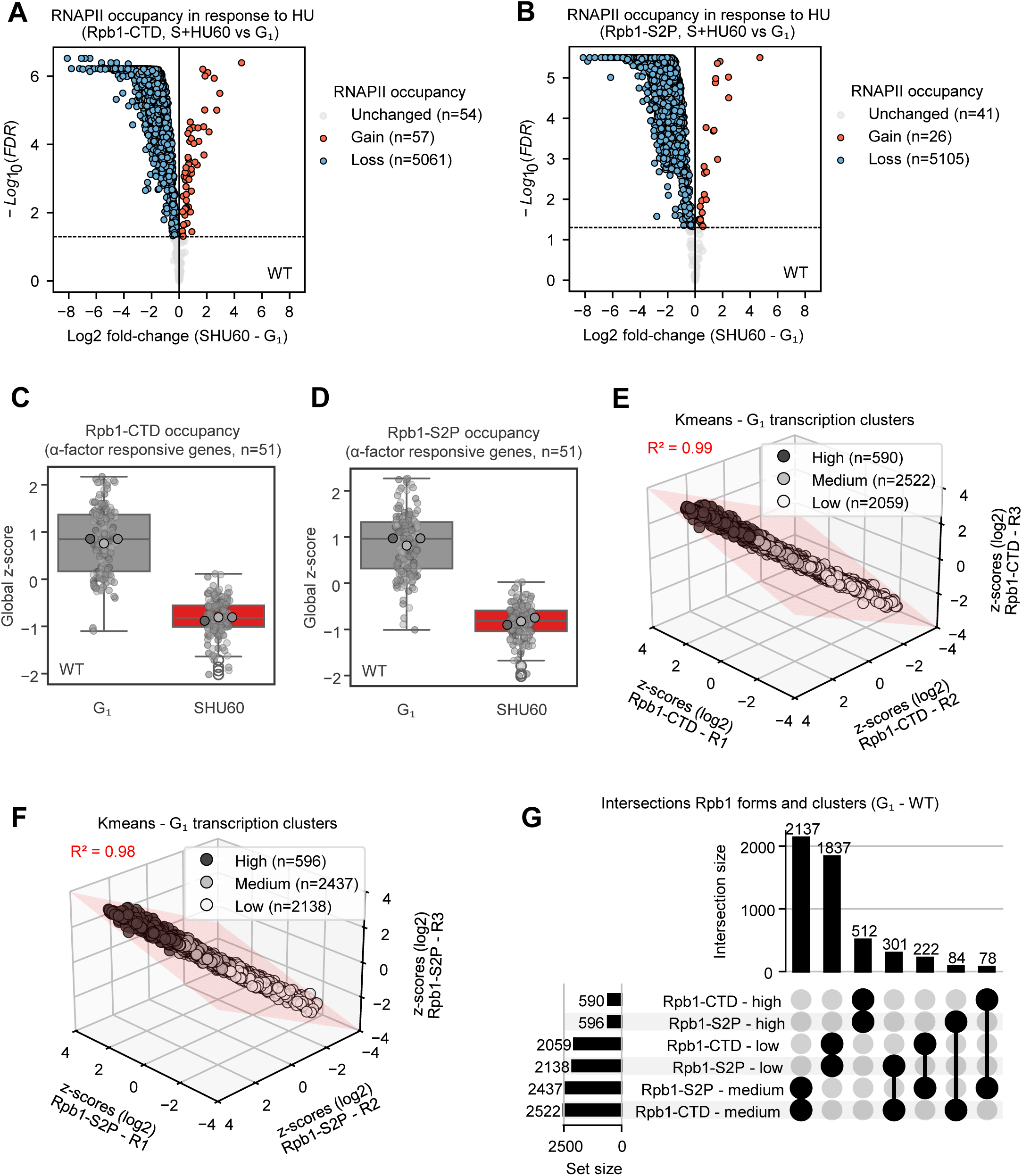
HU-induced replication stress induces RNAPII eviction from chromatin. (A, B) Volcano plots showing HU-induced changes in Rpb1-CTD (A) and Rpb1-S2P (B) chromatin binding in S+HU60 versus G₁ in the WT strain (Log2 fold change [S+HU60 vs G₁]). Colored dots indicate genes with significantly altered chromatin-binding scores (adjusted p-value < 0.05; Gain [Red, Log2FC > 0], Loss [Blue, Log2FC < 0]). (C, D) Boxplots of global z-scores for Rpb1-CTD (C) and Rpb1-S2P (D) occupancy at α-factor-responsive genes in G₁ (grey) compared to S+HU60 (red) in WT across three biological replicates. (E, F) 3D scatter plots showing G₁ transcription levels and replicate reproducibility ($n=3$) for Rpb1-CTD (E) and Rpb1-S2P (F) in the WT strain. Genes are grouped into High, Medium, and Low clusters via K-means clustering (k = 3) on log2-transformed, standardized RPKM scores. Planar regression coefficients (R^2^) are indicated. (G) Upset plot showing the intersections of gene clusters defined by Rpb1-CTD and Rpb1-S2P signal intensities (High, Medium, and Low expression clusters via K-means, k = 3) in the WT strain across three biological replicates.

**Figure S5:**
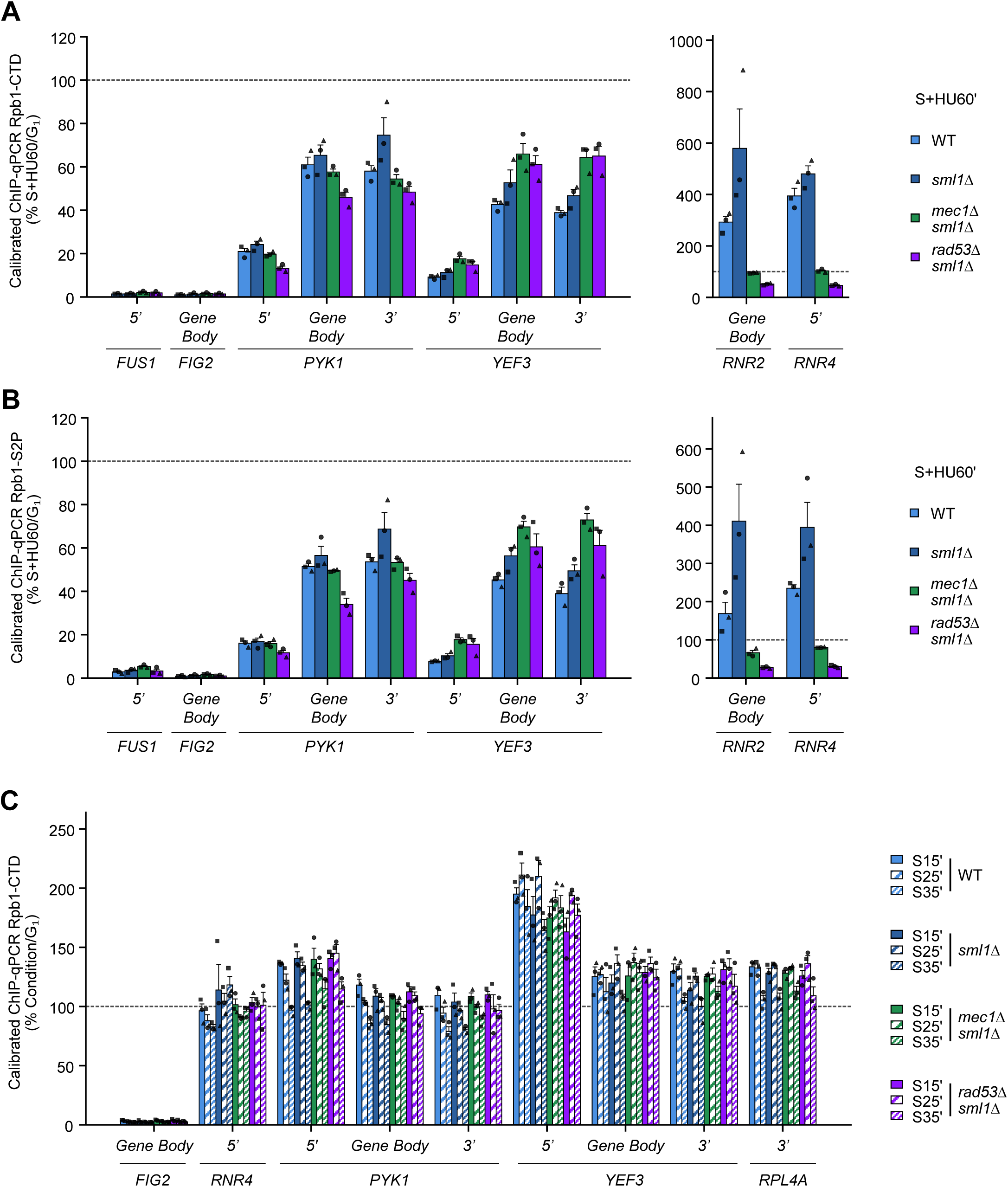
RNAPII dynamics during S-phase challenged or not by HU-induced replication stress. (A–C) Quantitative ChIP-qPCR analysis of RNAPII occupancy in Rpb1-CTD (A) and Rpb1-S2P (B) enrichment in S+HU60 relative to G₁. Enrichment was measured across the α-factor-responsive genes *FUS1* and *FIG2*, the glycolytic gene *PYK1*, the RiBi gene *YEF3*, and the stress-responsive genes *RNR2* and *RNR4*. (C) Rpb1-CTD enrichment during S-phase progression at 15, 25, and 35 min relative to G₁, measured across *FIG2*, *RNR4*, *PYK1*, *YEF3*, and the RP gene *RPL4A*. For all panels, values represent %Input normalized to the average %Input of *PYK1* and *PMA1* in *C. glabrata* and are expressed as a ratio relative to the G₁ condition. Individual biological replicates, together with the mean and SEM, are shown (N = 3).

**Figure S6:**
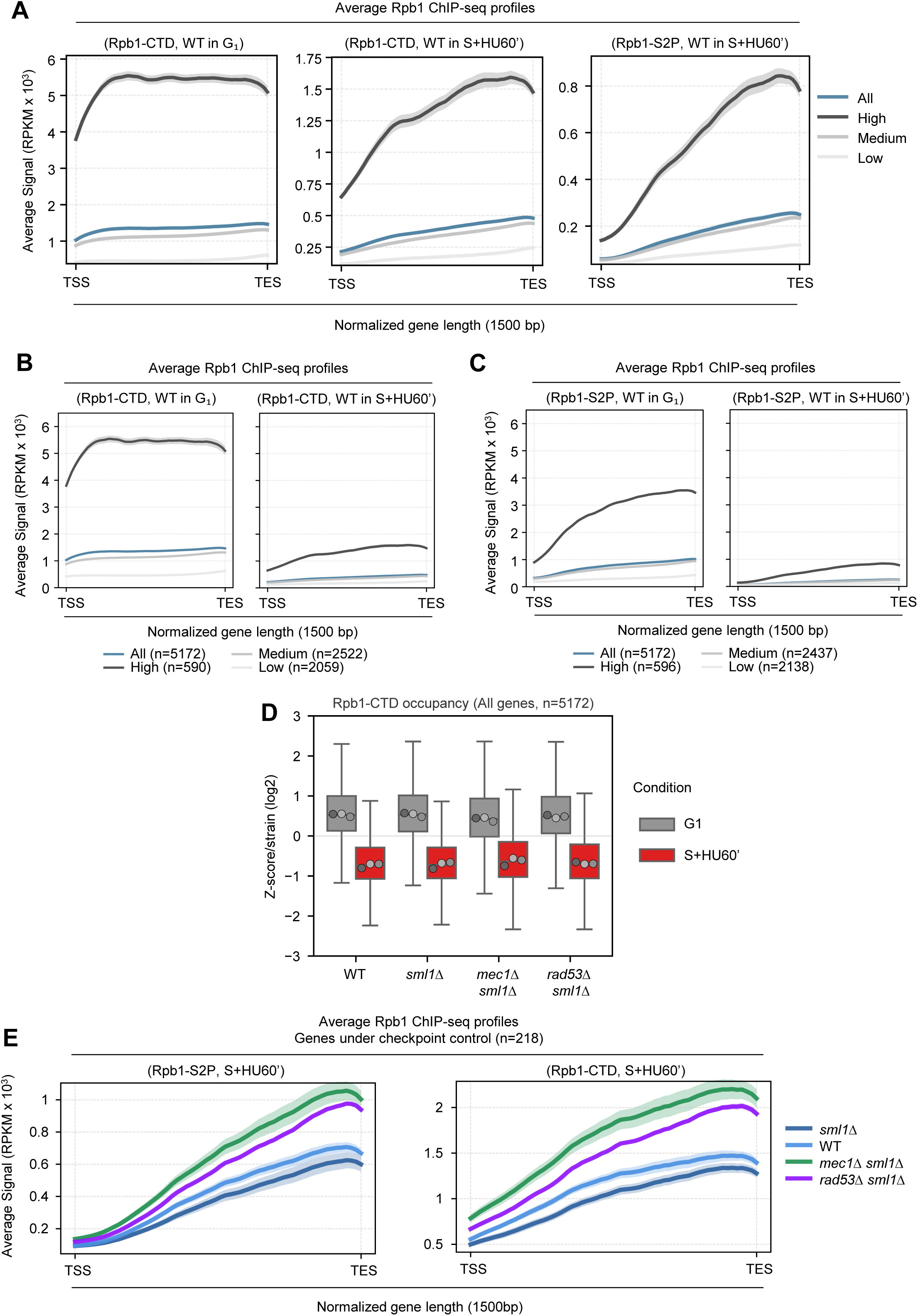
HU-induced replication stress shifts RNAPII Rpb1-CTD to Rpb1-S2P like profile. (A–C) Average Rpb1-CTD and Rpb1-S2P occupancy profiles from the TSS to the TES across three biological replicates. Data are expressed as spike-in-normalized RPKM across all RNAPII ORFs (n = 5172) or separated into High, Medium, and Low clusters, plotted over a normalized gene length of 1.5 kb. Mean profiles ± SEM are shown. (A) Profiles for Rpb1-CTD in WT cells in G₁ (1st panel) and S+HU60 (2nd panel), and for Rpb1-S2P in S+HU60 (3rd panel). For the Rpb1-CTD profiles, High, Medium, and Low clusters comprise 590, 2522, and 2059 genes, respectively; for Rpb1-S2P in S+HU60, they comprise 596, 2437, and 2138 genes, respectively. (B-C) Rpb1-CTD (B) or Rpb1-S2P (C) occupancy in WT cells in G₁ (1st panel) and S+HU60 (2nd panel). (D) Boxplot of Rpb1-CTD occupancy, expressed as a z-score per strain across all genes in G₁ (grey) compared to S+HU60 (red) across three biological replicates. (E) Average Rpb1-CTD and Rpb1-S2P occupancy profiles from the TSS to the TES for the S+HU60 condition across three biological replicates. Data are expressed as spike-in normalized RPKM for genes exhibiting Rpb1-CTD retention under hydroxyurea treatment that overlap between the *Mec1*-dependent and Rad53-dependent datasets, plotted over a normalized gene length (1500 bp). Mean profiles ± SEM are shown. (Left) Rpb1-CTD occupancy. (Right) Rpb1-S2P occupancy.

**Figure S7:**
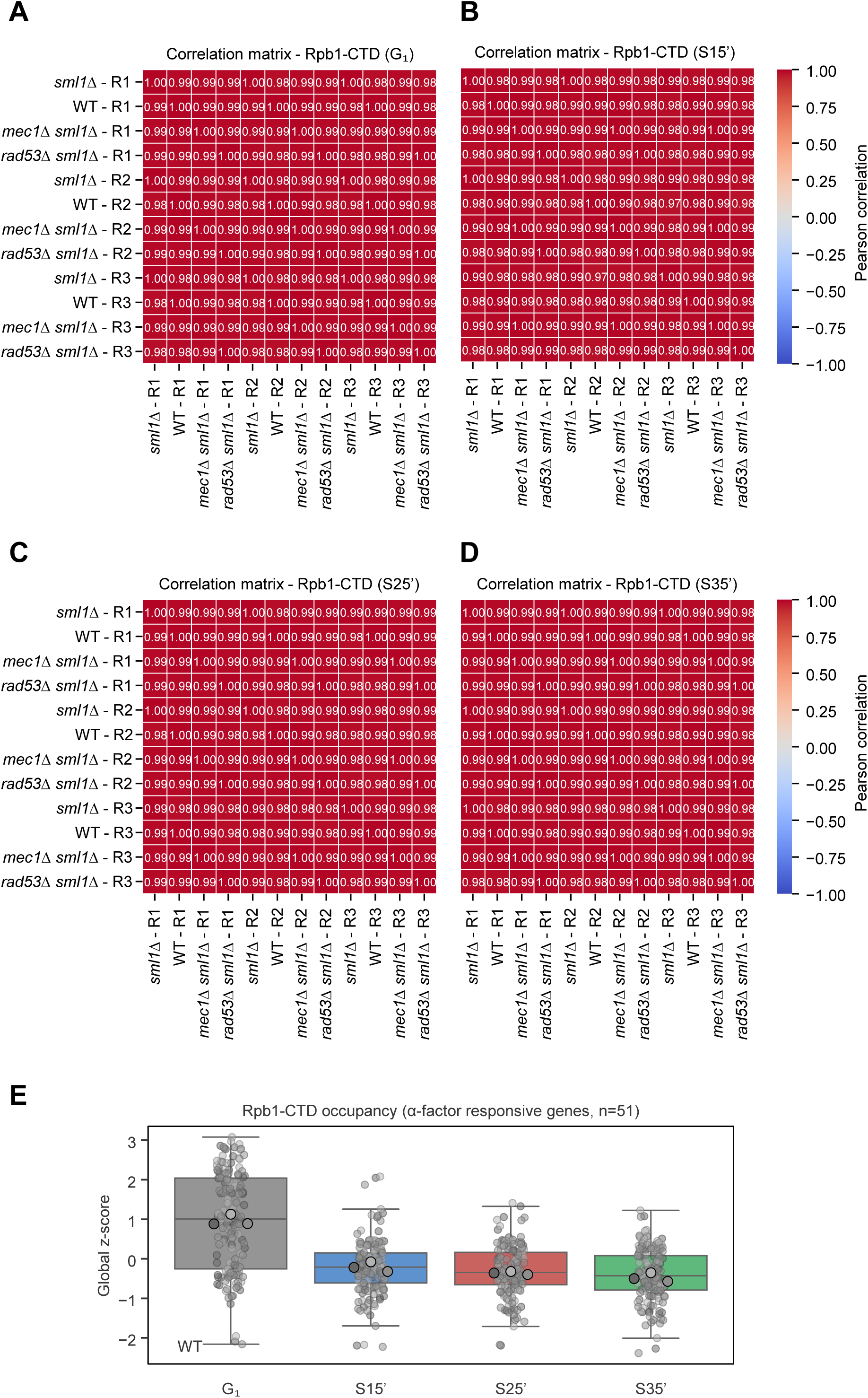
Rpb1-CTD correlation matrices for quantitative ChIP-seq in S-phase. (A-D) Pearson correlation heatmaps assessing biological reproducibility. Pairwise Pearson correlation coefficients (r) between replicate scores of Rpb1-CTD across strains and conditions in G₁ (A) and normal S-phase for 15 (B), 25 (C) and 35 (D) minutes. The color scale ranges from −1 to 1, indicating negative to positive correlation. N=3 biological replicates. (E) Boxplot of Rpb1-CTD occupancy, expressed as a global z-score at α-factor-responsive genes in G₁ (grey) and normal S-phase for 15 (blue), 25 (red) and 35 (green) minutes in the WT strain across three biological replicates.

**Figure S8:**
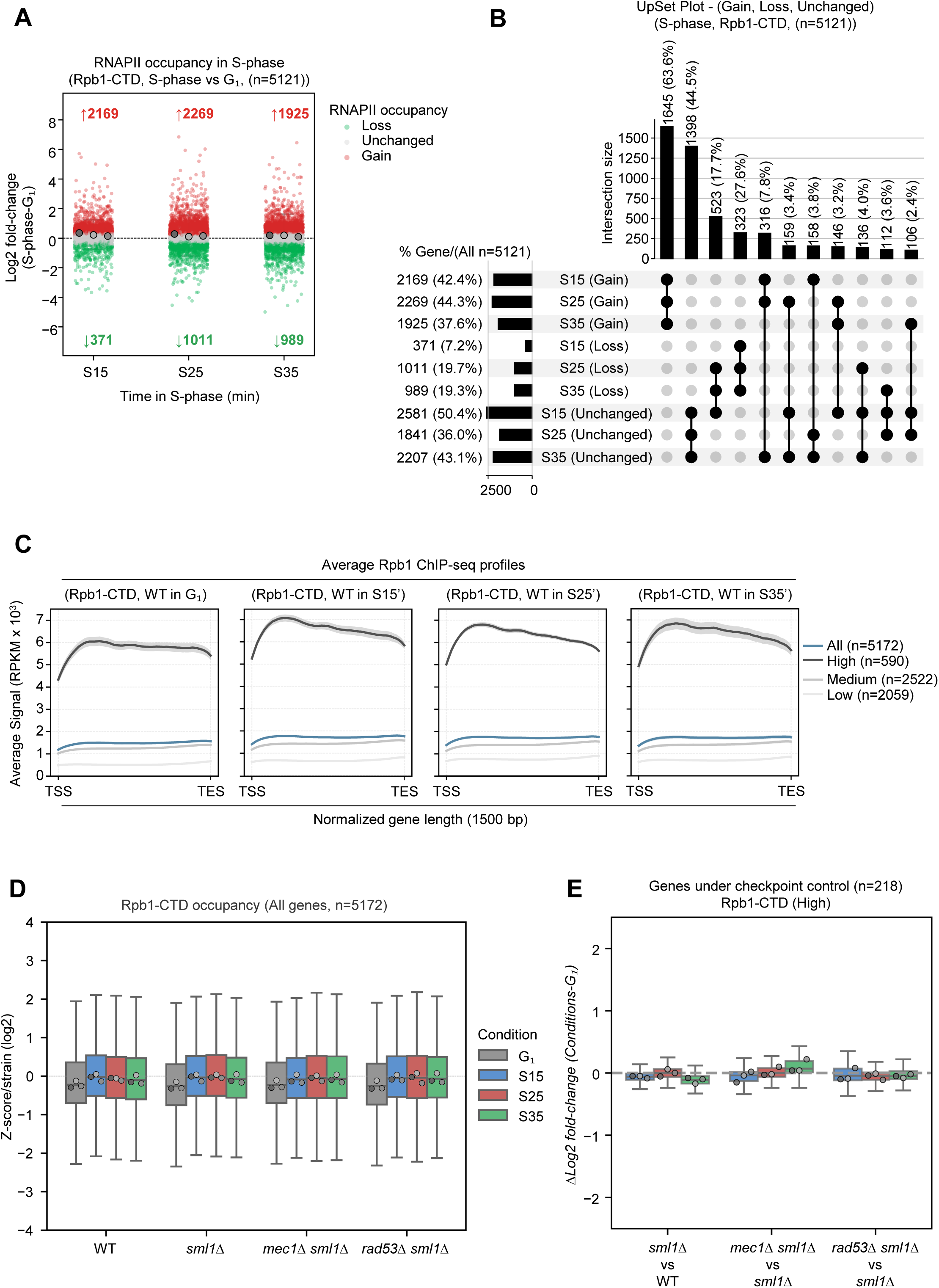
RNAPII eviction from chromatin is specific to replication stress. (A) Boxplot showing S-phase-induced changes in Rpb1-CTD chromatin binding in S-phase at 15, 25, and 35 minutes compared to G₁ in the WT strain (expressed as Log2-fold change [S vs G₁] across three biological replicates). Colored dots represent genes with a significantly different chromatin-binding score (adjusted p-value < 0.05; Log2-fold change > 0 [Gain, Red] or < 0 [Loss, Green]). (B) Upset plot showing the intersections of genes displaying Gain, Loss, or no change in Rpb1-CTD chromatin binding during S-phase (S15, S25, and S35) compared to G₁ in the WT strain across three biological replicates. Horizontal bars on the left indicate the percentage of genes in each individual category relative to the total number of genes, while vertical bars and intersecting points highlight the overlap of shared categories across the three time points. (C) Average Rpb1-CTD occupancy profiles from the TSS to the TES across three biological replicates. Data are expressed as spike-in normalized RPKM across all RNAPII ORFs or separated into High, Medium, and Low clusters, plotted over a normalized gene length (1500bp). Mean profiles ± SEM are shown for the WT strain: G₁ (1^st^ panel), S15 (2^nd^ panel), S25 (3^rd^ panel), and S35 (4^th^ panel). (D) Boxplot of Rpb1-CTD occupancy, expressed as a z-score per strain across all genes in G₁ (grey), S15’ (blue), S25’ (red) and S35’ (green) across three biological replicates. (E) Boxplot showing S-phase-induced changes in Rpb1-CTD chromatin binding in mutant strains relative to controls (expressed ΔLog2-fold change [S vs G₁]) for genes under checkpoint control under HU-replication stress. N = 3 biological replicates.

**Figure S9:**
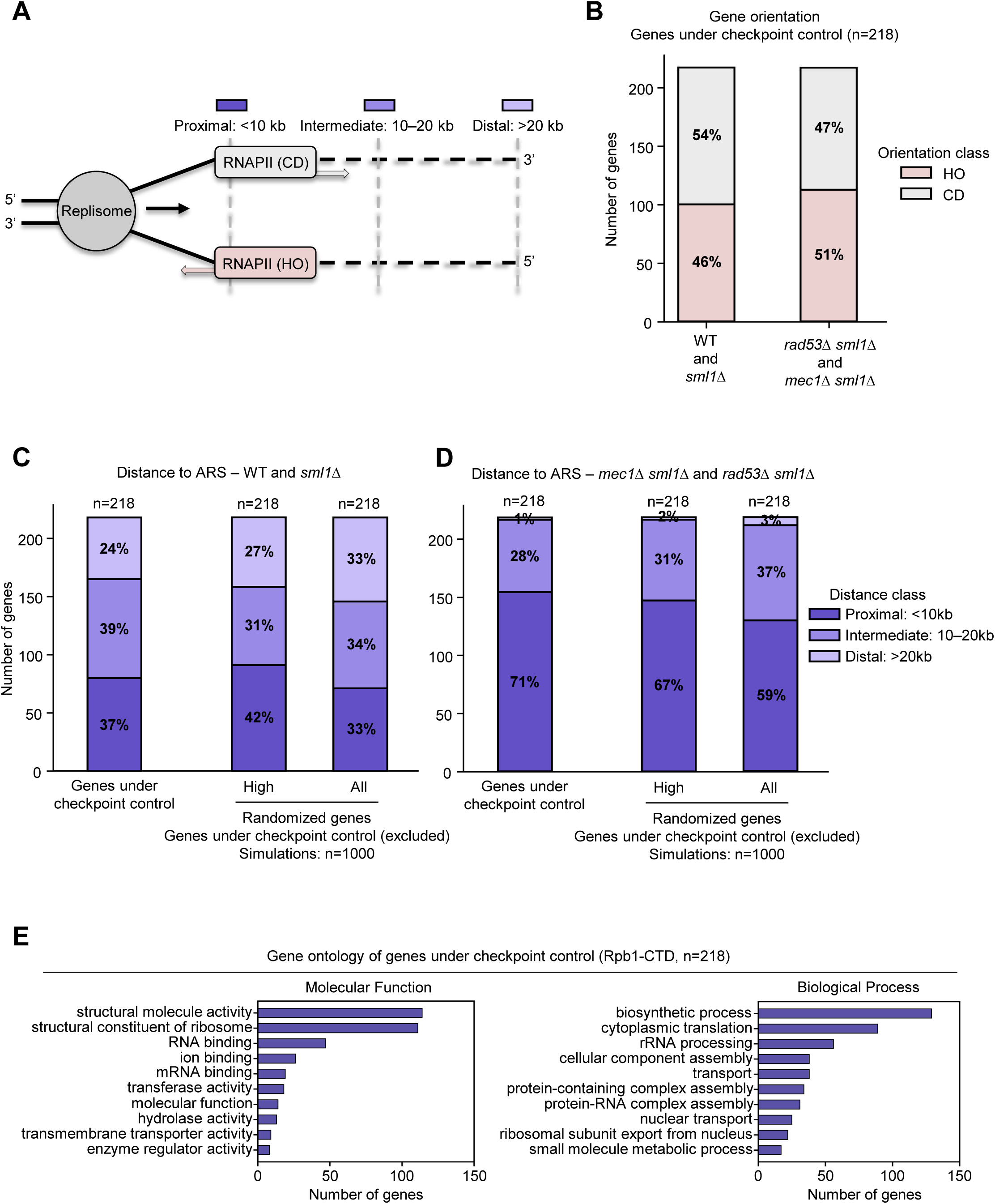
Genomic features of genes under checkpoint control during HU stress: orientation and proximity to active origins. (A) Schematic representation of gene orientation and genomic distance features used to assess gene orientation and proximity to replication forks. (B) Orientation of genes under checkpoint control, head-on (HO) or co-directional (CD) in respect to replication forks. Stacked bars display absolute gene numbers (y-axis) and percentages for orientation (HO vs. CD). (C, D) Distance of genes under checkpoint control to the nearest active ARS. Stacked bars display absolute gene numbers (y-axis) and distance proximity classes (<10kb, 10-20kb}, >20kb). (C) Control strains (WT and *sml1Δ*) replication patterns: Shows the orientation distribution for genes under checkpoint control, and the proximity distribution for an expression-matched randomized set of genes or a total yeast genome random set (excluding genes under checkpoint control) using a 1,000-iteration Monte Carlo simulation. (D) Checkpoint-deficient strains (*mec1Δ sm1Δ* and *rad53Δ sml1Δ*) replication patterns: Same distributions as in (C) evaluated under the *mec1Δ sm1Δ* and *rad53Δ sml1Δ* backgrounds. (E) Gene Ontology (GO) slim mapping of Biological Process/Molecular Function enrichment among genes exhibiting Rpb1-CTD retention under hydroxyurea treatment that overlap between the Mec1-dependent and Rad53-dependent datasets across three biological replicates.

**Figure S10:**
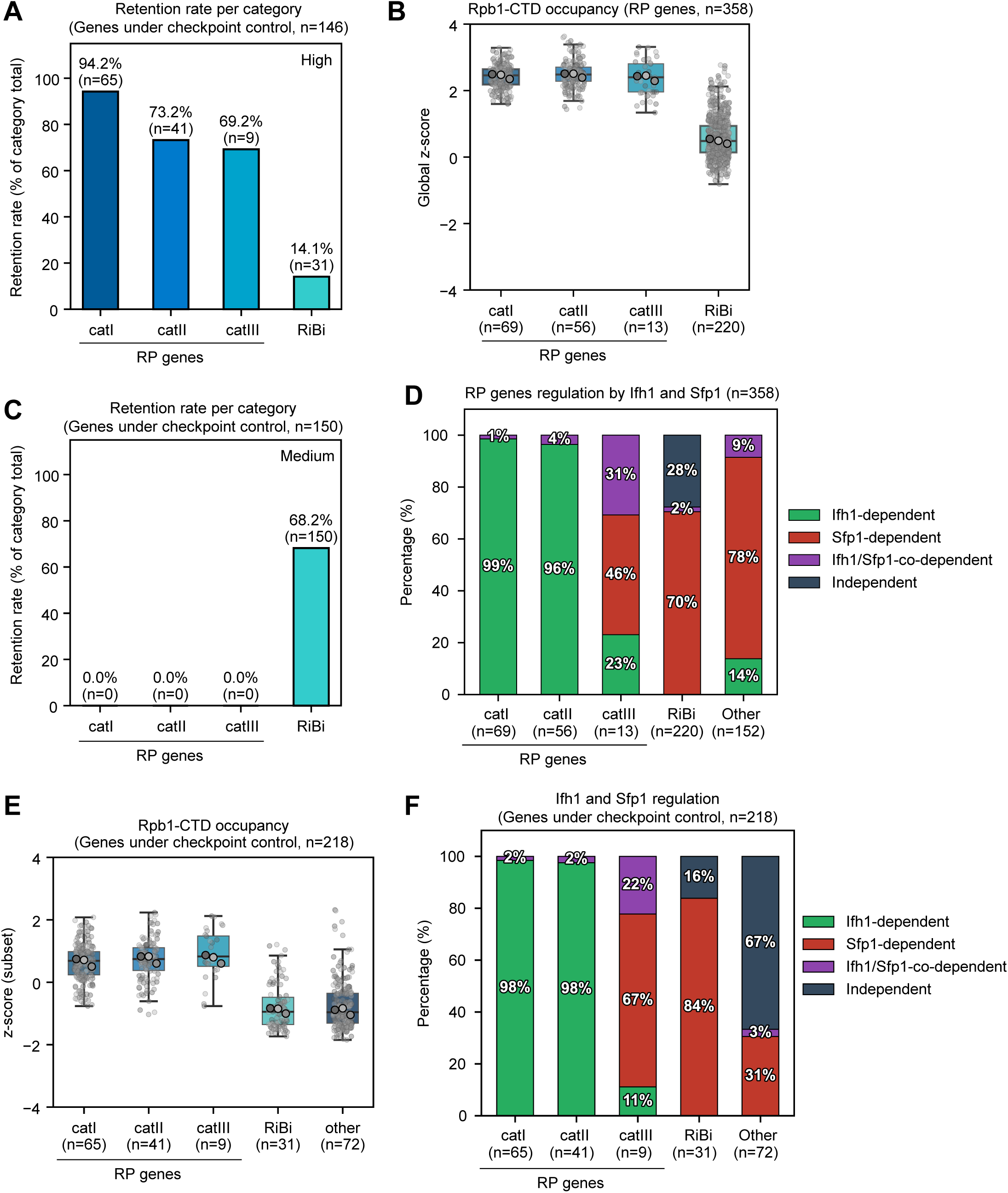
The majority of checkpoint-controlled genes are controlled by Ifh1 and Sfp1 transcription factors. (A, C) Bar plots showing the retention rate (% of retained genes over the total category) of RP classes and RiBi genes under checkpoint control in the High (A) and Medium (C) clusters. (B, E) Boxplots of global Rpb1-CTD occupancy z-scores across all genes (B) or restricted to checkpoint-controlled genes sorted by categories (E) (catI, catII, catIII, RiBi, and “other” representing non-RP checkpoint-controlled genes) in WT G₁ cells across three biological replicates. (D, F) Bar plots showing the percentage of genes dependent on Ifh1, Sfp1, or both across RP categories, RiBi genes, and “Other” genes for all analyzed genes (D) and checkpoint-controlled genes only (F). Transcription factor dependencies were determined by reanalyzing data from Zencir et al. 2020.

**Figure S11:**
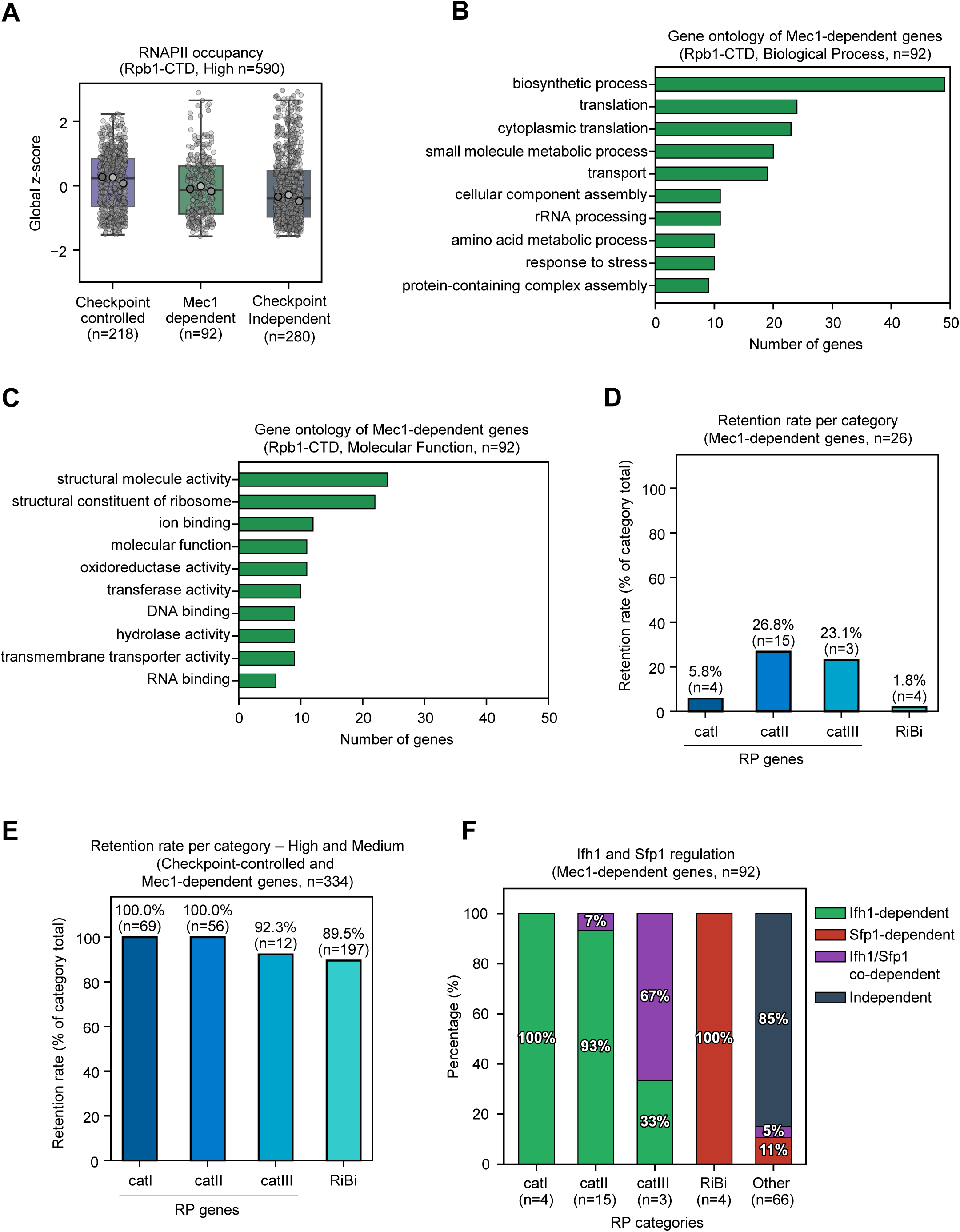
The majority of Mec1-dependent genes are controlled by Ifh1 and Sfp1 transcription factors. (A) Boxplot of global Rpb1-CTD occupancy z-scores across checkpoint-controlled, Mec1-dependent and checkpoint-independent genes in the high occupancy cluster, in WT G₁ cells (N= 3 biological replicates). (B, C) Gene Ontology (GO) slim mapping enrichment among genes exhibiting Rpb1-CTD retention under hydroxyurea treatment that are Mec1-dependent (N= 3 biological replicates). (B) Biological Process GO terms. (C) Molecular Function GO terms. (D, E) Bar plots showing the retention rate (% of retained genes over the total category) of RP and RiBi genes under checkpoint control and/or Mec1-dependent in the High (D) and High + Medium (E) clusters. (F) Bar plot showing the percentage of genes dependent on Ifh1, Sfp1, or both across RP categories, RiBi genes, and “Other” genes for Mec1-dependent genes. Transcription factor dependencies were determined by reanalyzing data from Zencir et al. 2020.

## Experimental procedure

### Yeast strains, culture conditions and drop assay

All strains used are listed in Table 1. For liquid cultures, yeast extract peptone medium was supplemented with 2% glucose unless otherwise stated. *MAT*a cells were synchronized in G_1_ by adding α-factor (8 μg/ml, GenScript) for 135 min at 25°C unless otherwise stated. Arrest without buds was monitored by phase microscopy. G_1_ blocked cells were released into S phase by addition of 75 µg/ml Pronase (Sigma, 53702) and were treated or not with the indicated dose of HU (CliniSciences, AG000WIJ). Drops assays were done with cells adjusted to 1 x 10^7^ cells/ml. 10-fold serial dilutions were spotted on YPD plates ± HU. For epitope tagging (tagging Rpo31 and Rpo31-FRB with 3xPK), *S. cerevisiae* and *C. glabrata* strains were generated via homologous recombination. *S. cerevisiae* clones were screened by PCR to verify correct integration, and positive clones were confirmed for protein expression by Western blot. Similarly, hundreds of *C. glabrata* clones were screened by PCR, and correct candidates were subsequently verified for protein expression by Western blot.

### Human cell culture and DNA spreading

Human osteocarcinoma U2OS cells (ATCC, HTB-86) were cultured at 37 °C in a 5% CO2 incubator in McCoy’s 5a medium supplemented with 10% fetal calf serum (FCS) and 100 U/mL penicillin/streptomycin (PS). Cells were treated with HU (CliniSciences, AG000WIJ), ATRi (Med Chem Express, Ceralasertib/AZD6738, HY-19323) and Triptolide (Med Chem Express, HY-32735). For DNA fiber spreading, subconfluent cells were sequentially labelled first with 10 µM 5-iodo-2’-deoxyuridine (IdU) and then with 100 µM 5-chloro-2’-deoxyuridine (CldU) for 30 min. One thousand cells were loaded onto a glass slide (StarFrost) and lysed with spreading buffer (200 mM Tris-HCl pH 7.5, 50 mM EDTA, 0.5% SDS) by gently stirring with a pipette tip. The slides were tilted slightly and the surface tension of the drops was disrupted with a pipette tip. The drops were allowed to run down the slides slowly, then air dried, fixed in 3:1 methanol:acetic acid for 10 min, and allowed to dry. Glass slides were processed for immunostaining with mouse anti-BrdU (347580, Becton Dickinson, 1:200) to detect IdU, rat anti-BrdU (Ab6326, Abcam, 1/300) to detect CldU, mouse anti-ssDNA (DSHB, AB_10805293, 1/200) antibodies and corresponding secondary antibodies conjugated to various Alexa Fluor dyes. Nascent DNA fibres were visualized by using immunofluorescence microscopy (Zeiss ApoTome). The acquired DNA fibre images were analysed by using Fiji.

### DNA combing

*S. cerevisiae* cells expressing codon-optimized TK and ENT1 were grown to 6 x 10^6^ cells/ml in YPD medium, synchronized in G1 and released into S-phase with 0.2 M HU. Cells were labeled with 400 µg/mL BrdU (B5002, Sigma) added 75 minutes post-release. Cells were harvested in 0.1% sodium azide, washed, and treated with Zymolyase buffer (50 mM phosphate buffer pH 7, 50 mM EDTA pH 8, 1.2 M Sorbitol, 10 mM DTT, 2.5 mg/mL Zymolyase 20T, 07663-91, Nacalai Tesque). Cells were embedded in 2% low-melting agarose (LMA, 11AGAL0050, MPBiomedicals) plugs (90 µL/plug), digested in Proteinase K buffer (125 mM EDTA pH 9.5, 1% Sarkosyl, 1 mg/mL Proteinase K) at 37 °C for 24 hours, and extensively washed. Plugs were equilibrated in MES buffer (1X MES pH 6, 100 mM NaCl) and melted at 67 °C. Genomic DNA was digested overnight with β-agarase (M0392L, NEB) at 42 °C. DNA was combed onto silanized coverslips (10490802, PolyAn) using an automated pulling machine (250 µm/s, Genomic Vision), baked at 60 °C for 2 hours, and mounted on glass slides. Coverslips were denatured with 1 M NaOH for 20-22 minutes, blocked, and incubated sequentially with primary antibodies against BrdU (347580, Becton Dickinson, 1:20) followed by fluorescent secondary antibody Alexa 546 (A-11030, Sigma, 1:50), and ssDNA (1:10) followed by fluorescent secondary antibody Alexa 647 (A-21241, Invitrogen, 1:50), with intermediate washes. Slides were mounted in Prolong Gold (P36930, Invitrogen). Images were acquired using a Zeiss Apotome microscope. Track lengths were measured using MetaMorph software, and data were plotted using Python, with 100 to 200 tracks analyzed per sample. To evaluate reproducibility and variability, SuperPlots were generated. Statistical significance was determined using a one-way ANOVA on replicate median values followed by Tukey’s multiple-comparisons test (ns, p > 0.05; *p < 0.05; **p < 0.001; ***p < 0.0001) across three biological replicates, as described by Lord et al. (2021).

### Recovery assay

Exponentially growing cells (4 mL) were treated for 2 hours with the indicated doses of HU +/-Cordycepin 60 µg/ml (C2689, TCI) or +/- Rapamycin (F6131, UBPBio) at the indicated dose. Following treatment, cells were pelleted, washed twice with an equal volume of sterile water at room temperature, and resuspended in 4 mL of sterile water. Cell concentrations were determined using a CASY apparatus (OMNI Life Science), and cultures were diluted to 2 x 10^6^ cells/ml. Three consecutive 10-fold serial dilutions were performed to yield a final concentration of 2 x 10^3^ cells/ml. For each condition, 100 µl (∼200 cells) of the final dilution were spread onto duplicate YPD plates using glass beads to establish technical replicates. Plates were incubated at 30°C for 2–3 days until colonies formed. Colonies were counted for both technical replicates, and mean values were calculated. Relative survival percentages were then determined by comparing the mean colony count of each condition to the wild-type control.

### Quantitative Chromatin Immunoprecipitation and Sequencing (ChIP-seq)

50 ml of cells were collected in G_1_ or at the indicated time after release into S phase ± addition of HU and immediately cross-linked with a final concentration of 1% formaldehyde (F8775, Sigma) for 15 minutes. Cross-linking was quenched with glycine (125 mM final concentration, G8898, Sigma) for 5 minutes, followed by two washes with ice-cold 1x TBS. In parallel, log-phase cultures of *Candida glabrata* were grown to 1 x 10^7^ cells/ml and subjected to the same cross-linking and quenching procedure. To avoid potential physiological effects of HU on *C. glabrata*, both yeasts were cross-linked separately prior to mixing. Cell pellets were resuspended in 50 ml of 1x TBS, and concentrations were determined using a CASY apparatus (OMNI Life Science). *S. cerevisiae* cells were adjusted to 1 x 10^7^ cells/ml, and *C. glabrata* cells were added as a spike-in at a 1:20 ratio. Cells were pelleted, resuspended in 800 µl of ice-cold 1x TBS, transferred to FastPrep 2mL tubes (No. 11432350, MP Biomedicals), and centrifuged at maximum speed for 1 minute. Supernatants were removed, and pellets were stored at −20°C until further processing. Cell pellets were resuspended in 520 µL of lysis buffer (50 mM HEPES-KOH pH 7.5, 140 mM NaCl, 1 mM EDTA, 1% Triton X-100, 0.1% Na-deoxycholate) supplemented with 1 mM PMSF (P7626, Sigma), protease inhibitors (cOmplete Tablet (1 for 50 ml lysis buffer), Roche, 04693116001), and Phosphatase Inhibitor Cocktail 3 (P0044, Sigma) when targeting phosphoproteins. Cells were lysed by bead-beating (MB400 U, Yasui Kikai, Osaka) using 200 µL of Zirconium beads. One-quarter of the lysate (130 µL) was recovered and sonicated in a microtube AFA Fiber Pre-Split Snap-Cap 6×16mm glass tube (520045, Covaris) adapted for a Covaris E220e sonicator using the following parameters: water level 6, peak power 175 W, duty factor 10%, cycles/burst 200, and time 420 seconds. Sheared chromatin was transferred to 1.5mL Eppendorf tubes, and the volume was adjusted to 300 µL using lysis buffer with protease inhibitors. Chromatin was cleared by centrifugation for 3 minutes at 8,000 rpm at 4°C. Aliquots were collected: 250 µL for immunoprecipitation (IP), 12.5 µL for input (whole cell extract, WCE) and sonication efficiency checks, and 10 µL for Western blot analysis. Prior to cell lysis, Dynabeads were washed three times and resuspended in 500 µL of PBS, 0.5% BSA, 0.1% Tween, then incubated with antibodies on a rotating wheel for at least 2 hours at 4°C. Specifically, 2.5 µL of anti-Rpb1-CTD (Clone 8WG16, BioLegend, 1 mg/mL) or 2.5 µL of anti-Rpb1-S2P (Clone 3E10, Abcam ab5095, 1 mg/mL) was incubated with 12.5 µL of Dynabeads Protein G (10004D, Invitrogen). Alternatively, 2.5 µL of anti-PK (Anti-V5 tag, BioRad, MCA1360G) was incubated with 12.5 µL of Dynabeads Protein A (10002D, Invitrogen). Antibody-coupled beads were washed once with 1 mL of PBS, 0.5% BSA, 0.1% Tween, resuspended in an equivalent initial volume of lysis buffer, and added to the chromatin extracts. Samples were incubated overnight on a rotating wheel at 4°C. Beads were collected on a magnetic rack and washed sequentially with cold solutions: twice with lysis buffer, twice with high-salt lysis buffer (50 mM HEPES-KOH pH 7.5, 360 mM NaCl, 1 mM EDTA, 1% Triton X-100, 0.1% Na-deoxycholate), twice with wash buffer (10 mM Tris-HCl pH 8, 0.25 M LiCl, 0.5% IGEPAL, 1 mM EDTA, 0.1% Na-deoxycholate), and once with TE buffer (10 mM Tris-HCl pH 8, 1 mM EDTA). Protein-DNA complexes were eluted from the beads using 120 µL of elution buffer (50 mM Tris-HCl pH 8, 10 mM EDTA, 1% SDS) for 12 minutes at 65°C. To reverse cross-links, 80 µL of TE with 0.1% SDS was added to the eluates, and 175 µL of TE with 0.1% SDS was added to WCE samples before overnight incubation at 65°C. Samples were treated with 130 µL of TE containing 400 µg of RNase A (∼20-40 mg/ml, R4642-1G, Sigma) for 2 hours at 37°C, followed by protein digestion with 10 µL of Proteinase K (20 mg/mL, Sigma, P6556) for 2 hours at 37°C. DNA was supplemented with 50 µL of 5 M LiCl and purified using two rounds of Phenol:Chloroform:Isoamyl Alcohol (25:24:1, P2069, Sigma) extraction. DNA was precipitated by adding 1/10 volume of sodium acetate (S2889, Sigma), 26 µg/mL glycogen (10901393001, Roche), and two volumes of 100% ethanol overnight at −20°C. Samples were centrifuged for 45 minutes at full speed at 4°C, washed with cold 70% ethanol, and centrifuged for an additional 5 minutes. Dried DNA pellets were resuspended in 30 µL of 1x TE. An aliquot of 10 µL (diluted to 300 µL) was used for qPCR analysis, while the remaining sample was reserved for library preparation. ChIP-seq DNA libraries were generated using the NEBNext Ultra II DNA Library Prep Kit for Illumina (E7645L, NEB) with 96 Unique Dual Index Primer Pairs (NEB), and subsequently sequenced.

### Quantitative ChIP-qPCR and Data Analysis

qPCR amplification was performed using a LightCycler 480 II instrument (Roche) in 384-well plates with the LightCycler 480 SYBR Green I Master (Roche, 04887352001), according to the manufacturer’s instructions. Primer specificity was first evaluated *in silico* by blasting primer sequences against the *S. cerevisiae* reference genome (sacCer3, April 2011 version). Only primers that hybridized uniquely to the *S. cerevisiae* genome and showed no cross-hybridization with the *C. glabrata* spike-in genome (CBS138 latest version) were selected. Experimentally, primers were designed to generate amplicons between 60 bp and 80 bp. Their specificity and amplification efficiency were tested using a serial dilution of genomic DNA (10, 1, 0.1, and 0.01 ng/µL). The thermal cycling program consisted of an initial denaturation at 95°C for 10 min, followed by 45 cycles of 95°C for 10 s, 60°C for 15 s, and 72°C for 15 s. Melting curve analysis (“Tm calling”) and quantification (“Abs Quant: 2nd Derivative Max”) were executed using the LightCycler 480 software (Roche). All validated primers exhibited a single sharp peak in melting curves and an amplification efficiency between 1.98 and 2.05, with an R^2^ ≥ 0.999. Primer efficiency (E) was calculated from the slope (B) of the standard curve plotting quantification cycles (C_q_) against the log of DNA template concentrations using the equation E = 10^(−1/B)^. For normalization, %Input values in *S. cerevisiae* were adjusted for each locus relative to the average %Input of *C. glabrata* reference genes *PYK1* (CAGL0M12034g) and *PMA1* (CAGL0A00495g) for RNAPII ChIP (Rpb1-CTD and Rpb1-S2P), and relative to *C. glabrata* tDNA genes *tI(AAU)2* (CAGL0I06468r) and *tF(GAA)2* (CAGL0E03377r) for RNAPIII ChIP (Rpo31-FRB-PK) samples. Because tDNA sequences are nearly 100% conserved between both yeasts, species-specific amplification was achieved by designing primer pairs with one primer within the tDNA and the other in the unique genomic flanking region, preventing cross-amplification with *S. cerevisiae*. Similarly, *PYK1* and *PMA1* primers showed no in silico cross-amplification with *S. cerevisiae* genomic DNA. All primers used for qPCR are listed in Table2.

### Quantitative ChIP-seq data processing

Paired-end sequencing reads were trimmed using Trim Galore (v0.6.7) and aligned to the *Saccharomyces cerevisiae* (sacCer3 without chrM, April 2011 version) reference genome using Bowtie2 (v2.4.4). Quantitative spike-in calibration was performed by aligning reads to the *Candida glabrata* (CBS138 without chrM, s05-m03-r32) genome, and PCR duplicates were removed using samtools (v1.13). Normalized coverage files (bigWig) and quantitative matrices were generated using bamCoverage and computeMatrix (deepTools v3.5.6) to evaluate RNAPII (Rpb1, Rpo31) occupancy across genomic features. Downstream analyses, including log2 transformations, K-means clustering (k=3), and empirical Bayes moderated t-tests (Benjamini-Hochberg FDR correction, p < 0.05), were conducted using custom Python scripts to identify significantly altered occupancy levels and gene clusters. GO Slim enrichment and spatial distribution analyses relative to Autonomous Replicating Sequences (ARS) were performed using SGD annotations and Monte Carlo-style resampling controls.

### Quantitative ChIP-seq data processing (Supp)

Paired-end sequencing (150 bp) was performed, and the flow cell was demultiplexed into FASTQ files by the sequencing provider. Raw reads were trimmed using Trim Galore! in paired-end mode with the adapters - a AGATCGGAAGAGCACACGTCTGAACTCCAGTCA for read 1 and - a2 AGATCGGAAGAGCGTCGTGTAGGGAAAGAGTGT for read 2.

Trimmed reads were first aligned against the Saccharomyces cerevisiae reference genome (sacCer3) using Bowtie2 with the following parameters: --very-sensitive --no-unal --no-mixed --no-discordant -k 2. The resulting BAM files were sorted by genomic coordinates. Concurrently, paired reads that failed to map or successfully mapped to sacCer3 were retrieved using the --un-conc-gz and --al-conc-gz parameters, respectively. To achieve quantitative calibration (spike-in normalization), unmapped and mapped reads from the sacCer3 alignment were subsequently aligned against the *Candida glabrata* (CBS138) genome using the same Bowtie2 parameters. Reads mapping to both genomes were filtered out to eliminate cross-species mapping artifacts. PCR duplicates were removed using samtools markdup. Final read counts were obtained using samtools stats on the filtered BAM files.

Occupancy ratios were determined using Input (WCE) and IP reads from both yeast species, as presented in Fig. S3A. Quantitative bigWig coverage files representing RPKM-normalized values were generated using bamCoverage (--binSize 25, --normalizeUsing RPKM, --extendReads), and the specific spike-in calibration scaling factor was applied to the IP samples using --scaleFactor.

Matrix-tab quantitative score files corresponding to RPKM values were subsequently generated from these bigWig files using computeMatrix scale-regions. For Rpb1-CTD and Rpb1-S2P, matrices were generated from the transcription start site (TSS) to the transcription end site (TES) of verified open reading frames (ORFs) from the SGD database (using a custom BED file), normalizing gene lengths to 1,500 bp (--regionBodyLength 1500, -bs 25) and sorting by mean value (--sortUsing mean). The same approach was applied for Rpo31-PK ChIP, using computeMatrix with a region spanning −300 bp to +300 bp relative to genes (-b 300 -a 300), a body length of 75 bp (--regionBodyLength 75), and a bin size of 25 (-bs 25). The resulting RPKM-based score files were then loaded into a custom Python script for downstream analysis.

For quantitative downstream analyses, matrix output files generated with computeMatrix were processed using a custom Python script. Gene annotations from a custom BED file were matched line-by-line with each matrix across all biological replicates. For each gene, total RPKM-based signal scores were quantified by summing the bin values across the region, and log2-transformed total scores were calculated. To assess biological reproducibility and data consistency, Pearson correlation matrices of Rpb1 and Rpo31-Pk replicate scores across conditions were then generated from these normalized scores. Score columns corresponding to each yeast strain, condition, and replicate were standardized using StandardScaler from scikit-learn, and pairwise Pearson correlation coefficients (r) were calculated and visualized as heatmaps with a fixed scale ranging from −1 to 1.

To define G_1_ transcription levels in the wild-type strain, genes were grouped into distinct clusters based on their Rpb1 or Rpo31 signal intensities. This was performed either by dynamically using K-means clustering (k=3) on log2-transformed, standardized RPKM total scores across biological replicates (z-scores calculated on log2 values). Replicate reproducibility and clustering distributions were evaluated using 3D scatter plots, accompanied by planar regression calculations to determine the coefficient of determination (R²).

To identify differentially occupied genes for Rpb1, moderated t-tests were performed using a custom Python implementation of the empirical Bayes variance-stabilizing approach (equivalent to the eBayes function in the R limma package). Gene-wise residual variances were pooled and squeezed toward a global median prior variance using fixed degrees of freedom to prevent false positives arising from unstable variance estimates in low-replicate groups. Log2 fold-changes and ΔLog2 fold-changes were calculated between the treatment and control groups. To avoid confounding results, α-factor-responsive and hydroxyurea-induced genes were excluded from the Log2 fold-change comparisons.

For Rpb1, raw p-values were subsequently adjusted for multiple testing using the Benjamini-Hochberg false discovery rate (FDR) procedure. Genes exhibiting an adjusted p-value < 0.05 were classified as undergoing significant changes in RNAPII occupancy (gain or loss) and were represented using volcano plots or boxplots.

For RNAPIII, statistical significance of differences in occupancy across yeast strains was evaluated using the non-parametric Mann-Whitney U test on gene-wise average log2 fold-changes. Pairwise comparisons were performed between strains, and p-values were categorized into significance *levels: p < 0.001 (***), p < 0.01 (**), p < 0.05 (*), and p >= 0.05 (ns, not significant*). Relative percentage changes for each mutant strain compared to the control strain were quantified using the mean log2 fold-change differences normalized to the absolute mean control value.

GO Slim mapping and enrichment analyses for the gene clusters were performed using significant genes identified in the given analysis. Briefly, a reference GO Slim annotation table for *S. cerevisiae*, obtained from the Saccharomyces Genome Database (SGD; go_slim_mapping.tab), was loaded to map input gene lists against standard GO Slim terms across the three main ontology aspects: Biological Process (P), Molecular Function (F), and Cellular Component (C). Subsetted annotations were aggregated to compute absolute gene counts (Gene_count) for a given GO term.

To analyze the genomic distribution of genes relative to replication origins, the distance from each gene to the nearest active Autonomous Replicating Sequence (ARS) was calculated for both wild-type and checkpoint mutant backgrounds. Genomic distances were determined using BED files containing ARS activation kinetics and checkpoint-regulated ARS annotations. Genes were classified into three distinct distance intervals based on their proximity to the closest active ARS: proximal (<10 kb), intermediate (10-20 kb) and distal (>20 kb). To determine whether the observed ARS distance distributions deviated significantly from random genomic expectations, a Monte Carlo-style resampling control was implemented. For each evaluated gene set of size n, a corresponding control set of genes was randomly sampled from the genome, matching the transcription class distribution of the test set while excluding the genes of interest. This simulation was repeated over 1,000 independent iterations. The resulting distance counts across all simulations were averaged to construct a stochastic background distribution, serving as a robust baseline for comparison against both WT and checkpoint ARS patterns.

## Data availability

Data have been deposited in the NCBI GEO under accession number GSE348626.

## Acknowledgements

We thank the Montpellier Ressources Imagerie (MRI) imaging facility. We thank Benoit Palancade for strains. J.H thanks the Ligue contre le Cancer for a PhD fellowship. This work was supported by the Agence Nationale pour la Recherche (ANR-22-CE12-0031-01, “SCooTR”) and IUF to J.P.

## Author contributions

Conceptualization: JH and JP. Investigation: JH, SG and JP. Methodology: JH and JP. Formal analysis: JH and JP. Bioinformatics: JH. Validation: JH and JP. Visualization: JH and JP. Funding acquisition, Supervision and Project administration: JP. Writing – original draft – JH and JP. Writing – review & editing: JH, PP and JP.

